# Phenotype switching of the mutation rate facilitates adaptive evolution

**DOI:** 10.1101/2023.02.06.527392

**Authors:** Gabriela Lobinska, Yitzhak Pilpel, Yoav Ram

## Abstract

The mutation rate plays an important role in adaptive evolution. It can be modified by mutator and anti-mutator alleles. Recent empirical evidence hints that the mutation rate may vary among genetically identical individuals: empirical evidence from bacteria suggests that the mutation rate can be affected by translation errors and expression noise in various proteins. Importantly, this non-genetic variation may be heritable via a transgenerational epigenetic mode of inheritance, giving rise to a mutator phenotype that is independent from mutator alleles. Here we investigate mathematically how the rate of adaptive evolution is affected by the rate of mutation rate phenotype switching. We model an asexual population with two mutation rate phenotypes, non-mutator and mutator. An offspring may switch from its parental phenotype to the other phenotype. We find that switching rates that correspond to so-far empirically described non-genetic systems of inheritance of the mutation rate lead to higher rates of adaptation on various fitness landscapes. These switching rates can maintain within the same individuals both a mutator phenotype and pre-existing mutations, a combination that facilitates adaptation. Moreover, non-genetic inheritance increases the proportion of mutators in the population, which in turn increases the probability of hitchhiking of the mutator phenotype with adaptive mutations. This in turns facilitates the acquisition of additional adaptive mutations. Our results rationalize recently observed noise in the expression of proteins that affect the mutation rate and suggest that non-genetic inheritance of this phenotype may facilitate evolutionary adaptive processes.

## Introduction

### Non-genetic inheritance of a mutator phenotype

Mutators—individuals in a living population with an above-average mutation rate—often arise spontaneously in during evolution [1]. Mutators may facilitate adaptation as they allow a faster exploration of the space of mutations. Yet, because there are more deleterious than beneficial mutations [2], mutators become associated with a higher mutational load compared to non-mutators [3], [4]. The mutator state is commonly thought to be stably genetically inherited over generations, as a result of, for example, inactivating mutations in DNA mismatch repair genes [1], [5], [6] or in DNA polymerase genes [7].

In contrast, the possibility of non-genetic inheritance of the mutation rate has received little attention. It has been suggested that transient mutators could arise, for example, as a result of a translation error in DNA repair proteins that could lead to a very strong mutator phenotype [8]. This mutator phenotype could be inherited for a few generations via cytoplasmic inheritance of faulty proteins, despite not being encoded in the genotype. In this scenario, an individual’s mutation rate can increase for a few generations, generating beneficial mutations without a long-term genetic commitment to an elevated mutation rate and the accumulation of deleterious mutations it entails. Indeed a quantitative analysis has been suggested that most adaptive mutations that appears in evolving population could be due to transient, rather than genetically inherited mutators [9].

Accumulating empirical evidence suggests that the mutator phenotype can be non-genetically inherited. Translation errors often occur in genes involved in DNA repair and replication [10], potentially realizing the scenario proposed by [8]. Moreover, a mechanism for epigenetic inheritance of the mutation rate in *Escherichia coli* has been described by [11]. This mechanism relies on cytoplasmic inheritance of Ada, a DNA repair protein induced under high pH. Each cell carries, on average, a single Ada molecule. Therefore, during cell division a substantial proportion of daughter cells inherit zero Ada molecules, and thus experience an elevated mutation rate. Hence, stochastic fluctuations in Ada copy number can affect the mutation rate. Furthermore, *Ada* is positively auto-regulated through a mechanism that can intensify the difference in its active copy number, and hence mutation rate, among genetically identical cells.

A more extreme case in which the mutation rate is not inherited, that is, there is no correlation between the parent and offspring mutation rates, was recently studied by [12]. These authors found that fluctuations in the mutation rate increase the population mean fitness by producing a subpopulation with below-average mutation rate when the population is well-adapted, and a subpopulation with above-average mutation rate when there is potential for further adaptation. In a subsequent work, it has been suggested that this variation in mutation rate could increase population evolvability [13].

### Fitness landscapes

Biological fitness landscapes are notoriously difficult to investigate due to their high dimensionality. Indeed, a DNA sequence of length *N* corresponds to 4^*N*^ genotypes, and a protein sequence of length *N* corresponds to 20^*N*^ possible proteins. Considering the average length of a gene, or that of a protein, it quickly becomes obvious that all possible sequences cannot be possibly surveyed empirically. Existing experimental studies of fitness landscapes have focused either on local landscapes around proteins [14], [15] or short DNA sequences [16]. Traditionally, fitness landscapes were described as containing many “valleys”, corresponding to low fitness genotypes, and “peaks”, corresponding to high fitness genotypes. Therefore, a central problem in evolutionary biology was to explain how fitness valleys can be crossed when natural selection acts to eliminate low-fitness intermediate genotypes [17]. A few examples of fitness valley crossing have been suggested [18], [19], along with strategies for their crossing such as capacitance [20], [21], partial robustness [22], stress-induced mutagenesis [23], or phenotypic variation [24]. Yet, due to the hyper-dimensionality of biological fitness landscapes, fitness valleys could actually rarely exist, and fitness landscapes may thus be more highly navigable than often appreciated [25]. Hence, evolutionary adaptation over rugged landscapes would be more akin to a diffusion problem in which adaptation is dependent on the time needed to find a mutational path that does not contain a fitness valley [26]. The empirical evidence for either an abundance or lack of fitness valleys is scarce, as observing fitness valley crossings poses technical difficulties due to the very low frequency of the low fitness intermediate genotype.

### The inheritance mode of the mutation rate

We suggest that the mode of inheritance of the mutation rate lies on a spectrum: at one end, mutator alleles that arise as rare mutations in genes involved in DNA repair or replication are inherited with very high fidelity, with little to no stochastic effects; on the other end of the spectrum, frequent and stochastic fluctuations in concentrations of proteins that affect the mutation rate may be of such magnitude that there is effectively no correlation between parent and offspring mutation rates; and in the middle of the spectrum, is a range that we define here to be “non-genetic modes of inheritance”, e.g. of the type that can be attained through cytoplasmic inheritance of proteins, or transgenerational epigenetic inheritance of fluctuations in protein concentrations. Such phenomena may produce a partially heritable mutator phenotype that is transmitted across several cellular generations at an intermediate fidelity. Another potential state is aneuploidy, which arises at rates much higher than genetic mutations, and may produce a significant mutator phenotype [27]. This spectrum of inheritance of the mutation rate is a special case of the adaptation spectrum of inheritance [28], which ranks modes of inheritance from high fidelity, such as genetic inheritance, to very transient, such as physiological changes.

### Overview

Here, we investigate multi-step adaptive evolution with different modes of inheritance of the mutation rate. We have developed a Wright-Fisher model with explicit inheritance of two mutation rate phenotypes: non-mutator and mutator. The main parameters of our model are the switching rate from the non-mutator to the mutator phenotype and the potentially different reverse-switching rate from the mutator to the non-mutator phenotype. We also examine several values of the non-mutator mutation rate and the fold-increase in the mutator’s mutation rate and explore several fitness landscapes.

We estimate the switching rates for three empirically described systems for non-genetic inheritance of the mutation rate: aneuploidy, the Ada protein in *E.coli*, and cytoplasmic inheritance of mistranslated proteins, and find that these estimates correspond to high adaptation rates on various landscapes.

We suggest that the combinations of switching rates that lead to high adaptation rates fulfil two conditions. First, the switching rate from non-mutator to mutator needs to be high enough to ensure a high frequency of mutators at the mutation-selection balance, and second, low enough so that the association between the mutator and the mutations it generates is maintained. In this case, individuals that already acquired a portion of the mutations needed towards a high fitness genotype are more likely to keep the mutator phenotype that is needed for the acquisition of the additional missing mutations. Moreover, additional combinations of values for the two switching rates lead to high adaptation rates on smooth landscapes (i.e., without “valleys” and only a single “peak”). These combinations increase the probability of hitchhiking of the mutator phenotype with beneficial mutations, thus facilitating the acquisition of additional beneficial mutations.

## Models and methods

### A general evolutionary model for genetic and non-genetic inheritance of mutation rate

We consider an asexual haploid population with non-overlapping generations and constant population size. We model the effects of mutation, phenotypic switching, natural selection, and genetic drift using a Wright-Fisher (WF) model [29]. The Wright-Fisher model is commonly used to model evolutionary biology. In this model, a population is described by a probability vector that contains the frequency of each genotype that appears in the population. This population vector is modified by iterating mutation-selection-drift steps. In the mutation step, the population vector is multiplied by a transition matrix that gives the probability of mutation between each pair of genotypes. In the selection step, the frequency of each genotype is multiplied by its relative fitness, and then the population vector is normalized so that it sums to one by dividing with the population mean fitness. In the drift step, stochasticity is introduced by randomly sampling the population vector of the next generation from a multinomial distribution characterize by the population vector of the current generation after the mutation and selection steps. One mutation-selection-drift cycle corresponds to a single generation of the population [29].

In our models, individuals are fully characterized by their mutation rate phenotype, either non-mutator (*m*) or mutator (*M*), and their “genome’s” genotype, which determines their fitness. The genotype is specified differently in each fitness landscape (see below). The combination of these two characteristics is termed pheno-genotype [30] to emphasize that it is defined through a combination of a non-genetic and a genetic component, namely the mutator phenotype and genotypic mutations. The frequency of individuals with mutator phenotype *z* and genome genotype *g* is denoted by *f_zg_*. In all cases we assume no recombination and, therefore, complete linkage.

Our model is similar to existing models for the study of genetic mutators, which are modifier loci that determine the mutation rate [23], [31], [32]. However, it is different from other models in that our switching rates can be orders of magnitude higher compared to the mutation rate that generates genetic mutators [8]. Moreover, studies on the evolution of genetic mutators often neglect back-mutations from mutators to non-mutators [31], [33], [34]. However, in our model, we consider non-genetic mechanisms for the inheritance of the mutation rate that may exhibit high rate of reversibility.

#### Mutation

The pheno-genotype frequencies after mutation, 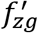, are given by

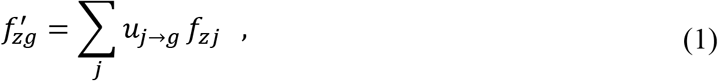

where *f_zj_* is the frequency before mutation, *j* is an index over all possible genotypes, *z* is the mutation rate phenotype, either *m* or *M* for non-mutator and mutator, respectively, and *u_j→g_* is the mutation transition probability from genotype *j* to genotype *g*. The specific transition probabilities *u_j→g_* in each fitness landscape are given below. In this mathematical notation *f_zg_* states the frequency of a pheno-genotype state whose mutation rate phenotype is *z* and whose genotype is *g*, where *f*’ represents the frequency after a mutation.

#### Phenotype switching

In every generation, an individual may switch its mutation rate phenotype from non-mutator to mutator with probability *γ*_1_ and or from mutator to non-mutator with probability *γ*_2_. We call these parameters “switching rates”. Thus, the pheno-genotype frequencies after phenotype switching are given by

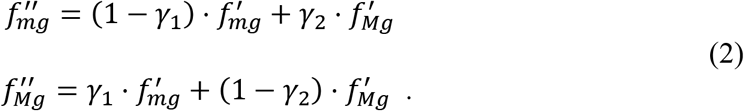

A schematic representation of the switching is shown in **Figure 1**. Later, we also extend our model to include a third transition pheno-genotype with a mutation rate that is intermediate between the non-mutator and the mutator mutation rates (see **Supplementary Figure 1**).

**Figure 1.**
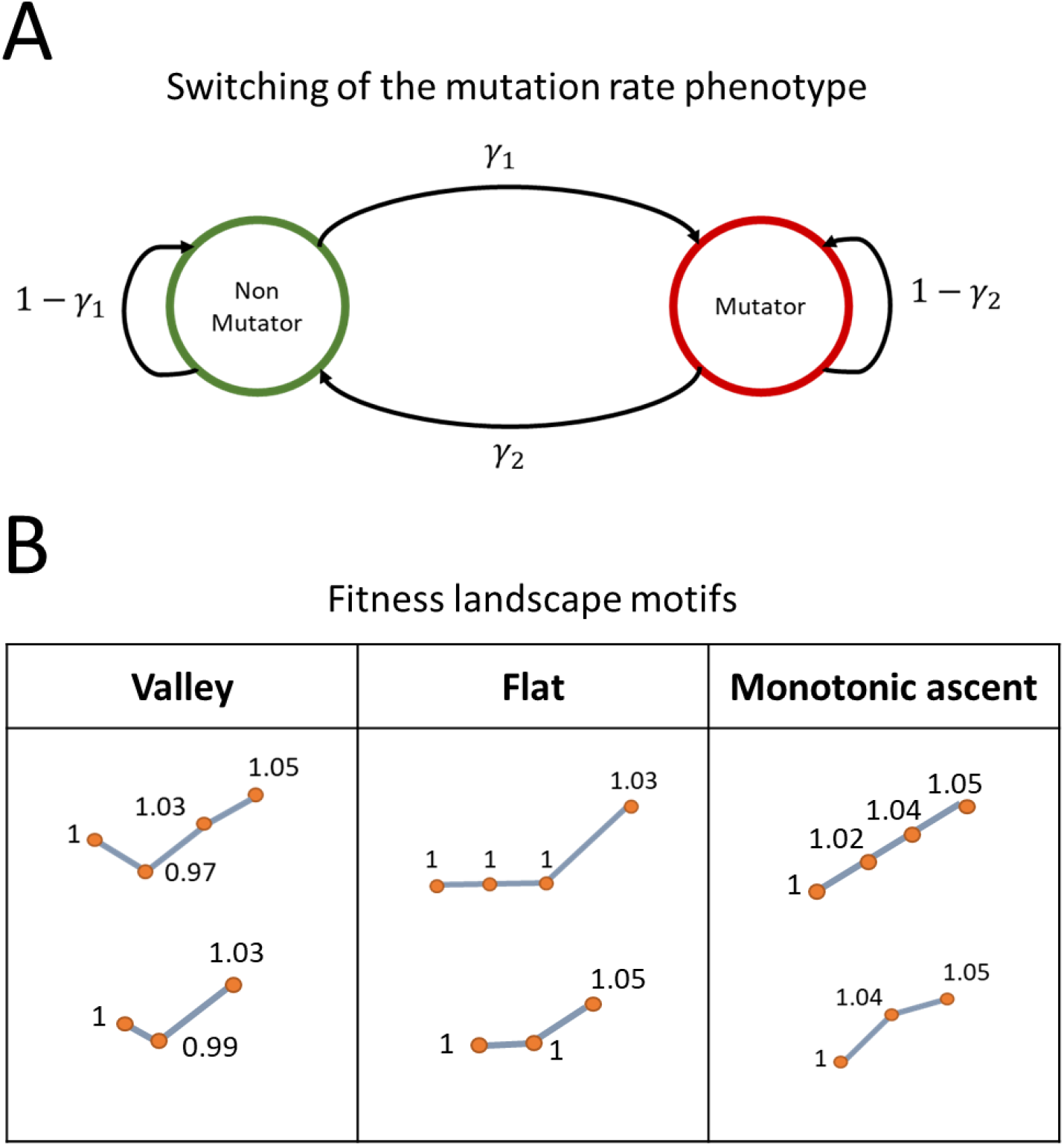
**(A) Switching between the non-mutator and the mutator phenotypes.** The switching rate from non-mutator to the mutator phenotype *γ*_1_ is the probability of an individual having a mutator phenotype given that its parent was a non-mutator. The switching rate from mutator to non-mutator *γ*_2_ is the probability of an individual having a non-mutator phenotype given that its parent was a mutator. **(B) Fitness landscape motif examples.** A fitness landscape motif is a succession of genotypes that are one mutation away from each other. The first genotype is considered the wild-type, and has a relative fitness of 1. The genotype with the highest number of mutations always has the highest fitness and is the adaptive genotype. Depending on the fitness of the intermediate genotypes, a fitness motif can be classified as valley, flat, or monotonic ascent.

#### Selection

In accordance with previous work on the evolution of modifiers, we assume that the mutator phenotype is neutral [33], [35], [36], that is, it does not directly affect fitness. In contrast to most previous work, we do not assume it is a gene, i.e. we do not assume genetic inheritance. Hence, we can focus on the fitness *w_g_* of genotype *g*, rather than the fitness of pheno-genotypes. The specific fitness values are determined by the fitness landscape, see below. The pheno-genotype frequencies after selection are given by

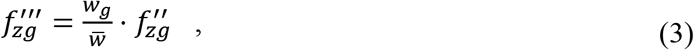

where 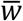 is the population mean fitness, 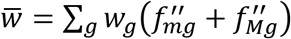.

#### Genetic drift

We model the effect of random genetic drift by drawing the number of individuals of each pheno-genotype from a multinomial distribution parameterized by the population size *N* and the pheno-genotype frequencies after mutation, phenotype switching, and selection, 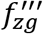.

#### Numerical analysis

The effects of random drift can sometimes be neglected (e.g., very large population size). In these cases, pheno-genotype frequencies are only affected by mutation, phenotype switching, and selection. We refer to this version of the model as the deterministic model and solve it numerically by iterating Eqs. (1–3) until the population reaches an equilibrium (that is, the frequencies of the pheno-genotypes do not change from generation to generation, 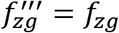). The equilibrium frequencies can also be obtained by numerically solving for the eigenvalues of the mutation-selection transition matrix, and normalizing the eigenvector corresponding to the leading eigenvalue. We observed that the values obtained by solving the eigenvalue problem corresponded closely to the values obtained from a numerical iteration of the model. The full model with drift is referred to as the stochastic model. It is simulated by iteration of Eqs. (1–3) and random sampling at each generation using a multinomial distribution.

### Fitness landscapes

#### Simple fitness landscape

To simplify our analysis before moving into biologically realistic fitness landscapes, we first study the rate of adaptation for a range of switching rates for a simple fitness landscape with a single adaptive peak.

##### Genotype

We consider genotypes with two or three major loci that affect fitness, in which beneficial mutations may occur, and a large number of background loci, in which only deleterious mutations may occur. Thus, the genotype is denoted by the number of mutations in the major loci and by the number of deleterious mutations accumulated in the background genomic loci. For example, the wild type genotype is *0\0*: it carries zero mutations in the major loci and zero mutations in the background loci, while a genotype *2\3* has two mutations in the major loci and three mutations in the background loci. The frequency of the genotype *0\0* is noted *f_0\0,0_* for non-mutators and *f_0\0,1_* for mutators (see above).

##### Fitness

We consider adaptation in two temporal phases. In the first phase, the wild-type genotype has the highest fitness. All other genotypes are maladapted, and hence the population converges to a mutation-selection balance around the wild-type genotype. During this phase, the fitness of the wild-type genotype *w*_0\0_ is set to 1. We assume that all mutant alleles in both the major and the background loci are deleterious with a selection coefficient *s* per mutation, such that the multiplicative fitness effect of *k* mutant alleles is (1 – *s*)^*k*^. For example, the fitness of the genotype 2\3 with two mutant alleles in the major loci and three mutant alleles in the background loci is *w*_2\3_ = (1 – *s*)^5^.

In the second phase, an environmental change occurs following which the fitness of the wild-type is no longer the highest. Rather, the double mutant in the major loci (when considering two major loci) or the triple mutant in the major loci (when considering three major loci) has the highest fitness. We still have *w*_0\0_ = 1. We define the vector (*w*_1\0_, *w*_2\0_) that describes the fitness values of individuals with one and two mutant alleles in the major loci and zero mutant alleles in the background loci; when considering three major loci, we use the vector (*w*_1\0_, *w*_2\0_, *w*_3\0_). Mutations in background loci still have a deleterious multiplicative effect. Thus, for example, the fitness of a genotype with two mutant alleles in the major loci and three in the background loci is *w*_2\3_ = *w*_2\0_ · (1 – *s*)^3^. The adaptive genotype is always the genotype with the largest number of mutant alleles in the major loci. Hence, *w*_2\0_ > *w*_1\0_ when considering two major loci and *w*_3\0_ > max(*w*_2\0_, *w*_1\0_) when considering three major loci.

##### Fitness landscape motifs

The fitness vectors (*w*_1\0_, *w*_2\0_) and (*w*_1\0_,*w*_2\0_, *w*_3\0_) allow us to introduce “fitness landscape motifs”. Fitness landscape motifs are a succession of genotypes neighbors in sequence space and their fitness. The wild-type fitness is *w*_0\0_. For example, if *w*_1\0_ < *w*_0\0_ < *w*_2\0_) when considering two major loci, the fitness motif will feature a fitness valley. If *w*_0\0_ < *w*_1\0_ < *w*_2\0_, the fitness motif is a monotonic ascent. Biological landscapes are composed of many such fitness landscape motifs, similarly to biological networks that feature network motifs [37]. We explore below how different combinations of switching rates influence adaptation over various fitness landscape motifs.

##### Mutation

The number of mutations per generation that occur at the background loci is Poisson distributed with expected value *U* or *τU* in individuals with non-mutator or mutator phenotype, respectively. Therefore, a genotype with *k* deleterious mutations will mutate to have *k+l* deleterious mutations with probability *e^−U^U^l^/l*! or *e^−τU^*(*τU*)^*l*^/*l*!. in individuals with non-mutator or mutator phenotype, respectively (*l* is assumed to be positive as we assume no back mutations). Mutations occur at the major loci with probability *μ* = *U/n* or *μ* = *τU/n*, according to the phenotype, where *n* is the total number of loci in the genome and *μ* is the per-locus mutation rate.

The number of mutations in the major loci is binomially distributed. The parameters of the binomial distribution are the number of major loci and the per locus mutation rate. We neglect the effects of back mutations, both in the numerical model and the stochastic model. This is because of the probability of a back-mutation following a mutation is negligible, as both of these events are rare. See Supplementary material for a formal description of the mutation transition probabilities.

### Complex and empirical fitness landscapes

We explore several fitness landscapes: NK landscapes with various ruggedness, commonly used to model biological fitness landscapes [38], [39], and an empirical landscape derived from the fungus *Aspergillus niger*, established by [40], [41]. A description of these landscapes and the choice of wild-type genotypes can be found in the Supplementary material.

### Values for model parameters

#### Mutation rates

The size of the *Saccharomyces cerevisiae* genome is 1.2 · 10^7^ bp [42], while the size of the *Escherichia coli* genome is about 5 · 10^6^ bp [43]. The proportion of deleterious mutations, out of all mutations, is about 40% [2], [44]. The mutation rate per bp for *S. cerevisiae* is about 10^−10^ per bp [45]; for *E.coli*, it is of the order of 10^−9^ – 10^−10^ per bp [46], [47]. Hence, we explore deleterious mutation rates from 4 · 10^−5^ to 10^−3^ per genome per cell cycle.

#### Fold-increase in the mutation rate in mutators

Most deletion mutants for genes involved in DNA repair in *S. cerevisiae* exhibit increase in mutation rate from 2- to 200-fold [48]. Mutator strains in *E. coli* exhibit a 50- to 1000-fold increase in mutation rate [49]. In this paper, we explore a fold increase in mutation rate for a mutator, *τ* = 10 and *τ* = 100, which is in the range of empirically observed values for the two microbes.

#### Mutation rate phenotype switching rates

The two main parameters of our model are the switching rate from non-mutator to mutator *γ*_1_ and the reverse switching rate from mutator to non-mutator *γ*_2_. As a special case the two rates may be identical. At least three potential systems for non-genetic inheritance of the mutation rate have been described in the literature: the Ada protein in *E.coli;* the cytoplasmic inheritance of mistranslated faulty proteins that affect mutation rate; and aneuploidy. For each of these, we estimate an approximate value for *γ*_1_ and *γ*_2_.

##### Ada protein in E. coli

The Ada protein is involved in DNA repair under high pH. The number of Ada molecules per cell is Poisson-distributed with an average of ~1. Cells with zero Ada molecules cannot efficiently trigger DNA repair under high pH and are hence they function as potential mutators.

On average, cells with zero Ada molecules produce one Ada molecule per cell cycle or generation [11]. We assume that the production of Ada molecules in a population of cells with zero Ada molecules follows a Poisson process with an rate of one molecule per generation. Thus, the probability of transitioning from mutator to non-mutator phenotype is *1-e*~0.63. We assume that stationary frequency of cells with zero Ada molecules is 25% (reported 20-30% in [11]). Solving for the probability of transition from cells with non-zero Ada molecules to cells with zero Ada molecules, we estimate this probability to be 0.21 (see Supplementary material, “Calculating the switching rate for Ada system”). Hence, for the Ada system of inheritance of the mutation rate, we have *γ*_1_ = 0.21 and *γ*_2_ = 0.63.

##### Aneuploidy

The state of aneuploidy—an abnormal number of chromosomes in the cell—has been shown to be associated with higher mutation rate [27]. The rate of aneuploidy in yeast has been estimated at 10^−4^ for whole-chromosome duplication and at 10^−5^ for whole-chromosome loss [50], [51]. Other estimates are 6.7 · 10^−6^ chromosome duplication events per generation and 3 · 10^−5^ chromosome loss events per generation [52]. Whole-chromosome loss in aneuploids is potentially faster than in euploids [27], [53]. Hence, we estimate *γ*_1_ = 10^−4^ and *γ*_2_ = 10^−5^.

##### Cytoplasmic inheritance of mistranslated proteins

We consider a hypothetical protein involved in DNA repair. We assume that this protein has the average length of a protein, is present in a single molecule in each cell, and is synthetized only when diluted out. In this section, we will compute the rate of transgenerational loss-of-function of that protein due to mistranslation and cytoplasmic inheritance. The assumption of a single molecule in each cell is made for simplicity, as it avoids the arbitrary decision of determining how many functional protein copies are needed for a non-mutator phenotype.

The rate of amino acid substitution in *E. coli* is about 10^−4^ to 10^−3^ per codon [10], [54]. The average length of a protein in bacteria is about 300 aa [55]. Following estimations from an empirical fitness landscape [14], we assume that about 10% of point mutations abolish the function of the protein. Hence, the switching rate from non-mutator to mutator, assuming a mistranslation rate of 5 · 10^−4^, is *γ*_1_ = 1 – (1 – 5 · 10^−4^)^30^ = 0.0149. The switching rate from mutator to non-mutator would be the dilution rate, that is 0.5, multiplied by the probability that the new cell will synthetize a functional protein. Hence, *γ*_2_ ~ 0.5.

In *S. cerevisiae*, the rate of mistranslation is 4 · 10^−5^ to 7 · 10^−4^ per codon [56] and the average length of a eukaryotic protein is about 500 aa [55]. Hence, assuming a mistranslation rate of 10^−4^ per codon, we have *γ*_1_ = 1 – (1 – 10^−4^)^50^ = 0.005. The switching rate from mutator to non-mutator is *γ*_2_ = 0.5 · (1 – 10^−4^)^50^ = 0.4975 ~ 0.5.

A summary of the estimates for the two switching rates and other model parameters is in **Table 1**. In this study, we use 10^−6^ as a low bound on genetic inheritance of the mutation rate. Because we consider two mutation rate phenotypes, a switching rate of 0.5 indicates that the mutation phenotype of the offspring is independent from the mutation rate phenotype of its parent. Switching rates higher than 0.5 signify that the offspring is more likely to have the mutation rate phenotype opposite of that of its parent than to have the same mutation rate phenotype.

**Table 1.** Summary of model parameters. Adapted from [23].

| Symbol | Name | Estimate | References |
| --- | --- | --- | --- |
| $N$ | Population size | $10^5$-$10^8$ | [72] |
| $U$ | Genome-wide mutation rate | 0.0004-0.003 | [73] |
| $n$ | Number of loci | 4,000-5,000 | [74] |
| $\mu$ | Per-locus mutation rate | $U/n$ | |
| $\tau$ | Fold-Increase in mutator mutation rate | 10-1,000 | [33], [75], [76] |
| $s$ | Selection coefficient against deleterious mutations | 0.001–0.03 | [2] |
| $H$ | Adaptation coefficient | 1–10 | [2] |
| $\gamma$ | Mutator phenotype switching rate | $10^{-6}$-0.5 | [8], [33] |
| $\delta$ | Switching rate from transition mutation rate phenotype | 0-1 | |

## Results

### Population at mutation-selection frequency balance

We first consider a population at the mutation-selection balance around the wild-type genotype. That is, at the mutation-selection balance the population is well-adapted to its environment and the frequencies of the pheno-genotypes do not change from generation to generation 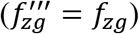 due to a balance between mutation and phenotype switching, which generate genetic and phenotypic variation, and selection and drift, which eliminate variation.

We focus on four pheno-genotypes (their frequencies appear in parentheses): non-mutators with wild type genotype (*m_0_*); non-mutators mutants (*m_1_*); mutators with wild type genotypes (*M_0_*) and mutators mutants (*M_1_*). Note that *f*_0\0,0_ = *m*_0_, *f*_1\0,0_ = *m*_1_, *f*_0\0,1_ = *M*_0_, and *f*_1\0,1_ = *M*_1_. Note that we consider the genotype 2\0 to be adaptive. Hence, the mutants 1\0,0 and 1\0,1 correspond to mutants that already have one out of the two mutations needed to become well-adapted. These four frequencies are plotted for several parameter sets in **Supplementary Figures 2–5.** Note that since the population is well-adapted to its environment, all mutants are deleterious. It is sufficient to focus on the four major genotypes due to our assumption on the population size (see **Supplementary Section B**), and following [57]. We also define *p_M_* as the frequency of mutators (as a sum of mutators with and without the mutation) in the population and *p_S_* the frequency of mutants (sum of mutator and non mutator mutants) in the population.

We consider two major loci, and an environmental change after which the genotype 2\0 is fitter than both 1\0 and 0\0. Adaptation is the appearance and subsequent fixation of the 2\0 genotype. We denote by *q* the probability of appearance and by *π* the probability of fixation conditional on appearance.

The probability that in the next generation no 2\0 mutant appears and is destined for future fixation in a population of size N is (1 – *qπ*)^*N*^. Assuming the number of generations until the appearance of a double mutant that goes to fixation is geometrically distributed, the adaptation rate *v* is defined as the probability that a single double mutant appears and escapes extinction by genetic drift,

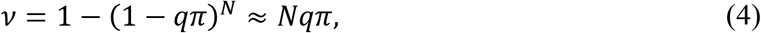

where the approximation holds when *Nqπ* is small. Note that the switching rate *γ*_1_ and *γ*_2_ affect the adaptation rate only indirectly due to its role in the MSB frequencies *m*_0_, *m*_1_, *M*_0_, and *M*_1_. We show in **Supplementary Section D**, that the probability of fixation of the double mutant given that it appeared, *π*, does not depend on the switching rate (see **Supplementary Figure 6**). Also, the time to fixation does not depend on the switching rate (see **Supplementary Figure 7**). Hence, in order to understand how the switching rates affect the rate of adaptation, we can focus on the probability of appearance of the double mutant.

The probability of appearance of the adaptive double mutant is derived from the frequencies *m*_0_, *m*_1_, *M*_0_, and *M*_1_. We neglect the frequency of a double mutant at the MSB (i.e., we disregard terms of order *μ^2^*) due to our assumption on the population size and the mutation rate (that is, 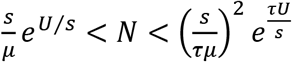, see **Supplementary Section B**). The probability *q* of appearance of double mutants as a result of a mutation in an existing single mutant in a population that currently does not have any double mutants is the sum of the probabilities of its appearance either from a mutator or a non-mutator

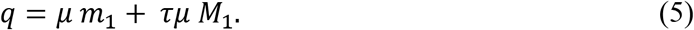

Here, *q* is dominated by the term *τμ M*_1_, i.e. appearance from a mutator (**Supplementary Figure 8**) and therefore maximization of *M*_1_ will also lead to maximization of the adaptation rate. Hence, the frequency *M*_1_ of the mutator mutant is of crucial importance to the rate of adaptation.

**Figure 2** shows the probability of appearance *q* depending on the two switching rates, *γ*_1_ and *γ*_2_. The three biological systems of non-genetic inheritance of the mutation rate, namely the fluctuation in amount of the Ada protein, translation error, and aneuploidy, are each marked as coloured stars according to their estimated switching rates. The probability of appearance is very high for *γ*_1_ > *τU*/2 when *γ*_2_ < *τU*/2, and for *γ*_1_ > *γ*_2_ when *γ*_2_ > *τU*/2. Otherwise, it is very low. The probability of appearance for additional parameter sets is shown in **Supplementary Figure 9**. We notice that all three systems of non-genetic inheritance of the mutation rate correspond to a high probability of appearance. The rate of adaptation for gradual switching of the phenotype are shown in **Supplementary Figure 10** and **11**. There are no major differences with **Figure 2**. We performed a sensitivity analysis for the number of loci *n* and found that it has a very limited effect on the relative rates of adaptation for various combinations of switching rates (see **Supplementary Figure 12**).

**Figure 2.**
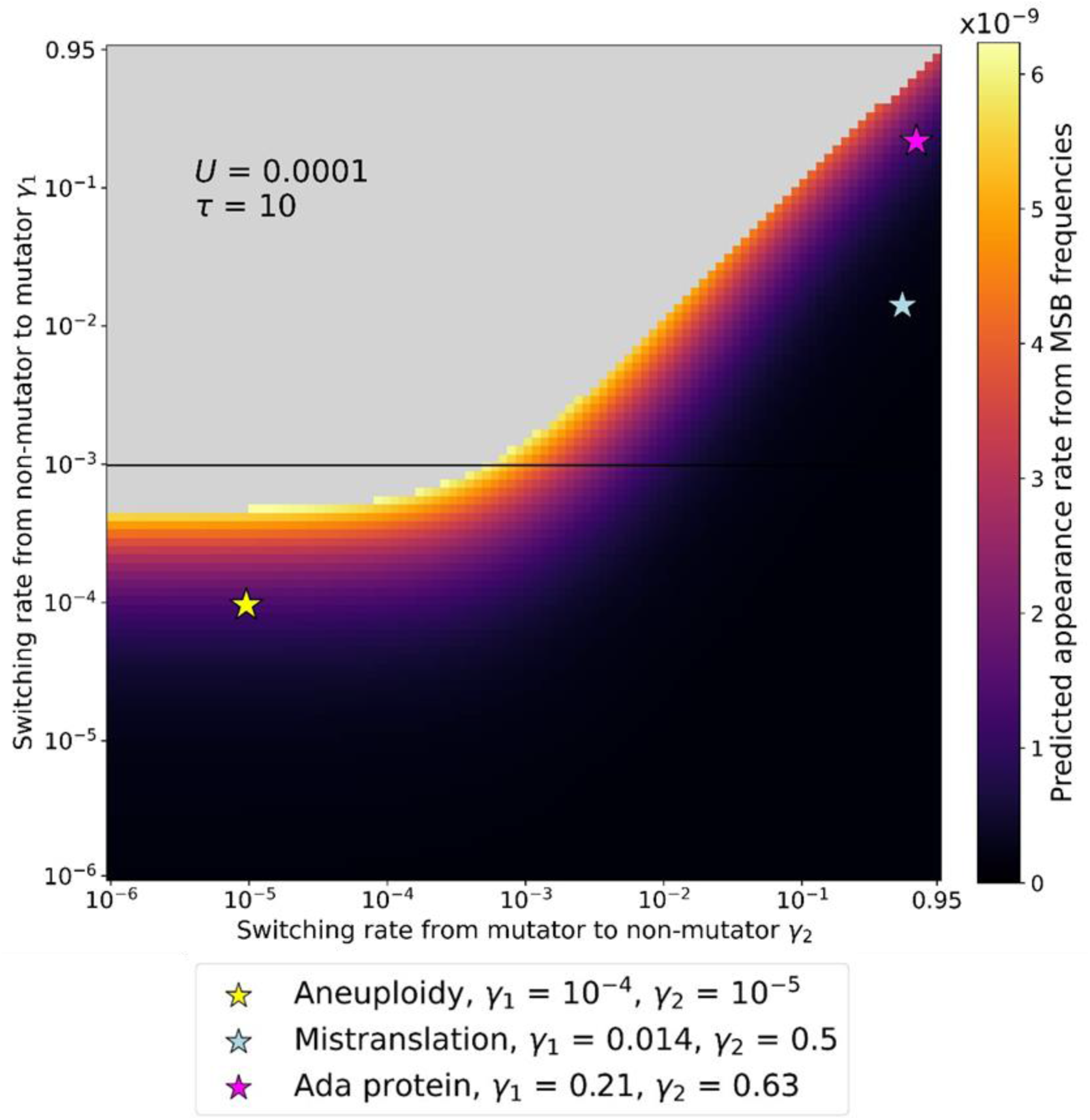
Adaptation is faster with intermediate switching rates. The adaptation rate based on mutation-selection balance (MSB) frequencies of mutators and mutants (Eq. 6). We assume that the double mutant is adaptive after an environmental change, and that the single mutant is deleterious. Grey area represents parameter combinations that give a MSB mutator frequency higher than 0.5. Coloured stars correspond to three systems of non-genetic inheritance of the mutation. The highest adaptation rates (bright yellow) occur for an intermediate rate switching to mutator, *γ*_1_ ≈ 10^−3^ (y-axis) and a wide range of switching back to non-mutator, 10^−5^ < *γ*_2_ < 10^−2^. Parameters: *s* = 0.03, *β* = 5000.

We also plotted the frequency of mutators *p_M_* and mutator mutants *M*_1_ in **Supplementary Figures 13** and **14**. When *γ*_1_ > *τU*/2 and *γ*_2_ < *τU*/2, and when *γ*_1_ > *γ*_2_ and *γ*_2_ > *τU*/2, we have *p_M_* ~ 1, which is biologically unrealistic: it means that the whole population has an elevated mutation rate. The frequency of *M*_1_ is correlated with the probability of appearance, which is consistent with the fact that the term *M*_1_ dominates the expression for *q* (see Eq. 5). We notice in particular that the frequency of mutator mutants is lower for *γ*_1_ > 10^−2^ and *γ*_2_ > 10^−2^, as expected from the observed rates of adaptation in this region of the parameter space.

Therefore, we postulate two factors that can increase the frequency of *M*_1_ at the mutation-selection balance, and therefore the adaptation rate: (i) because mutators are more likely than non-mutators to generate mutations, any effect that will increase the proportion of mutators *p_M_* at mutation-selection balance will increase the frequency of mutant mutators *M*_1_; (ii) maintaining the association between mutators and the mutants it generates, after they are generated. Below we explore each of these two effects.

#### Frequency of mutators at mutation-selection balance

We approximate the frequency of mutators at the mutation-selection balance (MSB). We denote by 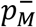 the frequency of mutators at MSB when *γ*_2_ is low and by 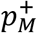 the frequency of mutators at MSB when *γ*_2_ is high.

Here, we follow [33], which studies the frequency of genetic mutators at the mutation-selection balance. Therefore, they assumed low *γ*_1_ and neglected *γ*_2_ altogether. They found that

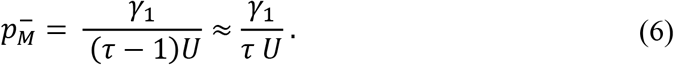

Eq. 4 provides a good approximation of the frequency of mutators *p_M_* at the mutation-selection balance when *γ*_2_ is low. When *γ*_2_ increases, the approximation becomes worse (**Supplementary Figure 15**). The maximal 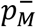 value is 0.5 (beyond that value, the mutator is no longer the minority phenotype), yielding *γ*_1_ = *τU*/2. Above *γ*_2_ > *τU*/2, the balance between the two mutation rate phenotypes is no longer expressed by Eq. 4.

When both switching rates are high, *γ*_1_, *γ*_2_ > *τU*/2, the indirect selection against the mutator phenotype becomes inefficient. Thus, the mutation-selection balance between the non-mutator and the mutator phenotypes can be approximated by the stationary distribution of a Markov chain, such that 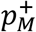, the frequency of mutators with high switching, is

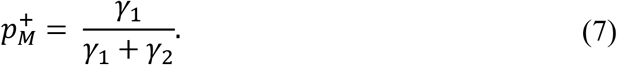

The frequency of mutators 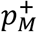 is maximised when *γ*_1_ = *γ*_2_.

Hence, the frequency of mutators *p_M_* is maximised at 0.5 by *γ*_1_ = *τU*/2 when *γ*_2_ < *τU*/2, and *γ*_1_ = *γ*_2_ when *γ*_2_ > *τU*/2 (**Supplementary Figure 13**).

#### Association between mutator and mutant phenotypes

Maximising the frequency of mutators is not sufficient to maximise the frequency of mutator mutants.

The frequency of the mutator phenotype is not independent from the mutations it generates: a large fraction of mutations are generated by mutators [8], [33], and inheritance of the mutator phenotype preserves the association between mutators with mutants.

The association between the mutator and the mutant phenotypes can be quantified by

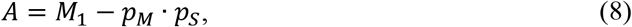

which is defined as the association between the mutator phenotype and the single mutant genotypes minus their expected frequency that would be expected if they were independent, that is given by their frequency product. Thus, if the mutator and mutant state were independent then *A*=0. The association between mutator and mutant phenotypes is plotted for several parameter sets in **Supplementary Figure 16**.

When *γ*_2_ < *τU*/2, we observe that association increases with *γ*_1_, reaching its peak at around *γ*_1_ = *τU*. When *γ*_1_ > *τU*, the whole population has the mutator phenotype, and the association decreases to 0, since *M*_1_ = *p_S_* and *p_M_* = 1. For *γ*_2_ > *τU*/2, we have high association for *γ*_1_ = *γ*_2_.

However, we also observe that for very high *γ*_1_ and *γ*_2_, the frequency of *M*_1_ decreases. This is because when *γ*_1_ = *γ*_2_ = 0.5, there is effectively no inheritance of the mutator state, and *A* = 0. Hence the frequency of mutator mutants *M*_1_ is not maximized. The threshold for when inheritance becomes negligible seems to be about 10^−2^, as read from **Figure 2**.

We can therefore conclude that the rate of adaptation is maximised when the frequency of mutators *p_M_* is maximized, while keeping the association between mutators and mutants *A* high through maintaining a high inheritance of the mutator phenotype. These conditions are fulfilled when *γ*_1_ = *τU*/2 for *γ*_2_ < *τU*/2, and *γ*_1_ = *γ*_2_ < 10^−2^ for *γ*_2_ > *τU*/2. These conditions are met by the three biological systems of non-genetic inheritance described in the empirical literature (coloured stars in **Figure 2**) when *τ* = 10.

### Adaptation rate for different landscape motifs

In the previous section, we computed the rate of adaptation assuming two major loci, and the fitness of each of the two single mutants in a major locus is lower than that of the double mutant and wild-type. However, other landscapes are possible. In this section, we use simulations to estimate the rate of adaptation for several other landscapes with two major loci or three major loci.

We consider three main categories of fitness landscape motifs: (i) the “fitness valley”, where one of the intermediate genotypes has lower fitness than both wild-type and the adaptive genotype; (ii) the “flat” where all intermediate genotypes have the same fitness as the wild-type; and (iii) the “monotonic ascent” where each additional mutation results in an increase in fitness. Within each category, the relative values of the fitness of the intermediate genotypes are possible. The motifs considered in this paper are shown in **Figure 1B**, along with their names.

For each fitness landscape, we ran 1,000 simulations of the stochastic model for 45,000 generations (for two major loci) or 100,000 generations (three major loci). The frequencies of each pheno-genotype at generation 0 are the frequencies corresponding to the mutation-selection balance frequencies. In **Figure 3**, we plot the proportion of simulations in which the adaptive genotype has reached >99% in frequency after a given number of generations. Results for additional fitness landscape motifs are shown in **Supplementary Figure 17** and **18**. We observe that the highest rates of adaptation corresponded to the region with highest frequencies of mutator mutants at mutation-selection balance identified in the previous section. For landscapes featuring an increase in fitness with each subsequent mutation, high rates of adaptation were also observed when both *γ*_1_ < *τU*/2 and *γ*_2_ < *τU*/2. This cannot be explained by a high frequency of mutator mutants (see **Figure 2**). Rather, the high adaptation rate must have been due to the dynamics of the mutator frequency during evolution from the wild-type to the adaptive double or triple mutant.

**Figure 3.**
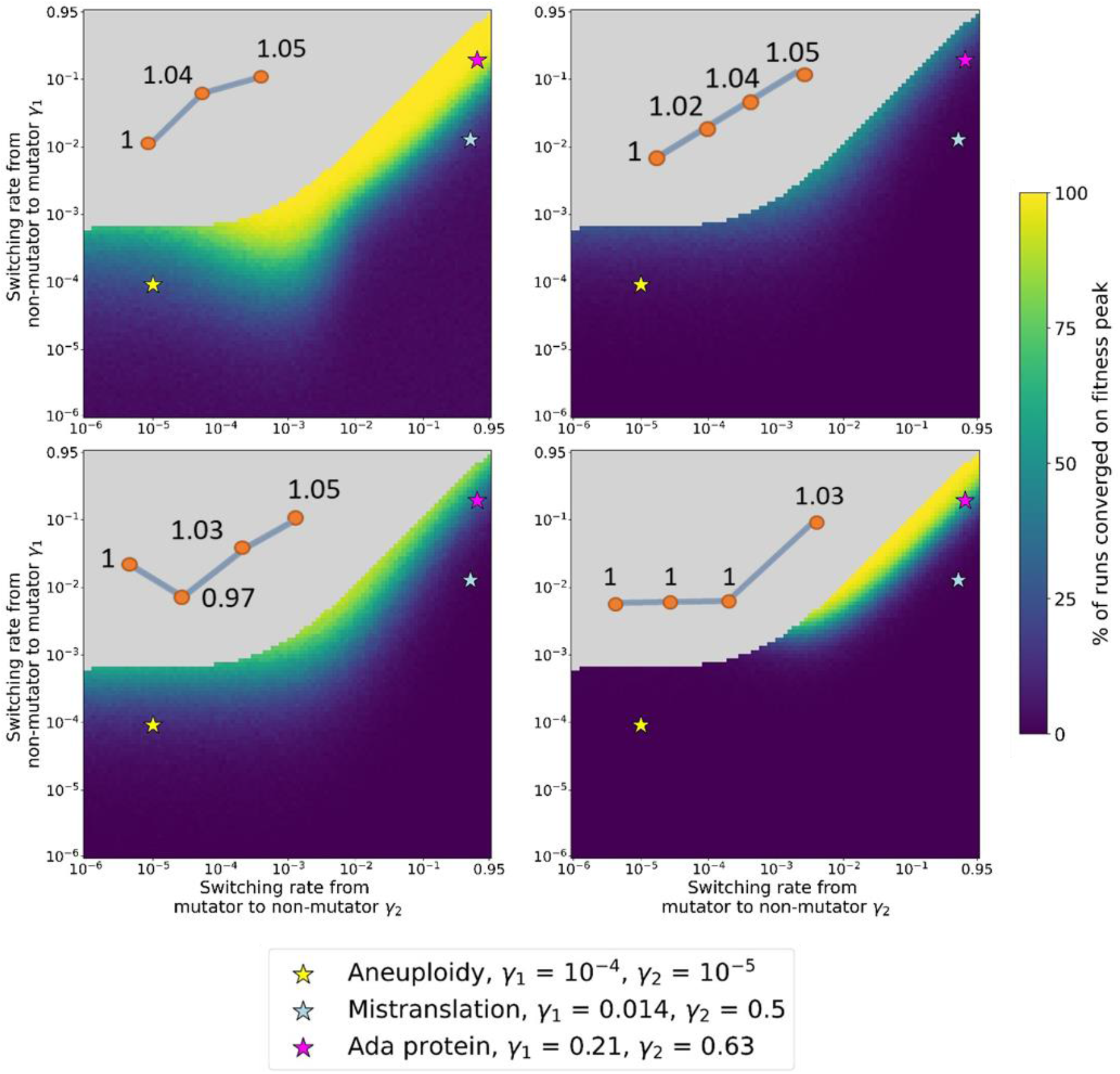
Dynamics of adaptation on different landscape motifs. The figure shows the proportion of populations that reached the adaptive genotype in simulations over different fitness landscapes. For progression after different numbers of generation, see Supplementary Figures 17 and 18. Cartoons show the fitness landscape shape, with numbers representing the relative fitness of sequential genotypes. Coloured stars represent three systems of non-genetic inheritance of the mutation rate. Parameters: *U* = 4 · 10^−5^, *τ* = 100, *s* = 0.03, *β* = 5000.

We hypothesized the following mechanism: the mutator being more likely to generate the genotypes that are intermediate between the wild-type and the adaptive genotype, it could hitchhike with one of the intermediate genotypes (either single mutants when considering two major loci, or double mutants when considering three major loci) that have higher fitness compared to the wild-type and increase in frequency. This increase in the frequency of intermediate genotypes with a mutator phenotype facilitates the appearance of genotypes with additional mutations, leading to a high rate of adaptation when both *γ*_1_ < *τU*/2 and *γ*_2_ < *τU*/2.

To test this hypothesis, we measured the mutator frequency during adaptive evolution (**Figure 4**). When the landscape motif features an increase in fitness with each additional mutation, the three mutations tend to occur and fixate very close together in time, and each fixation leads to an increase in mutator frequency (see bottom line, panel B with *γ*_1_ = *γ*_2_ = 10^−4^). The mutator frequency does not return to its mutation-selection balance frequency before the appearance of another adaptive mutant, hence the mutator frequency remains high, facilitating the generation additional mutations.

**Figure 4.**
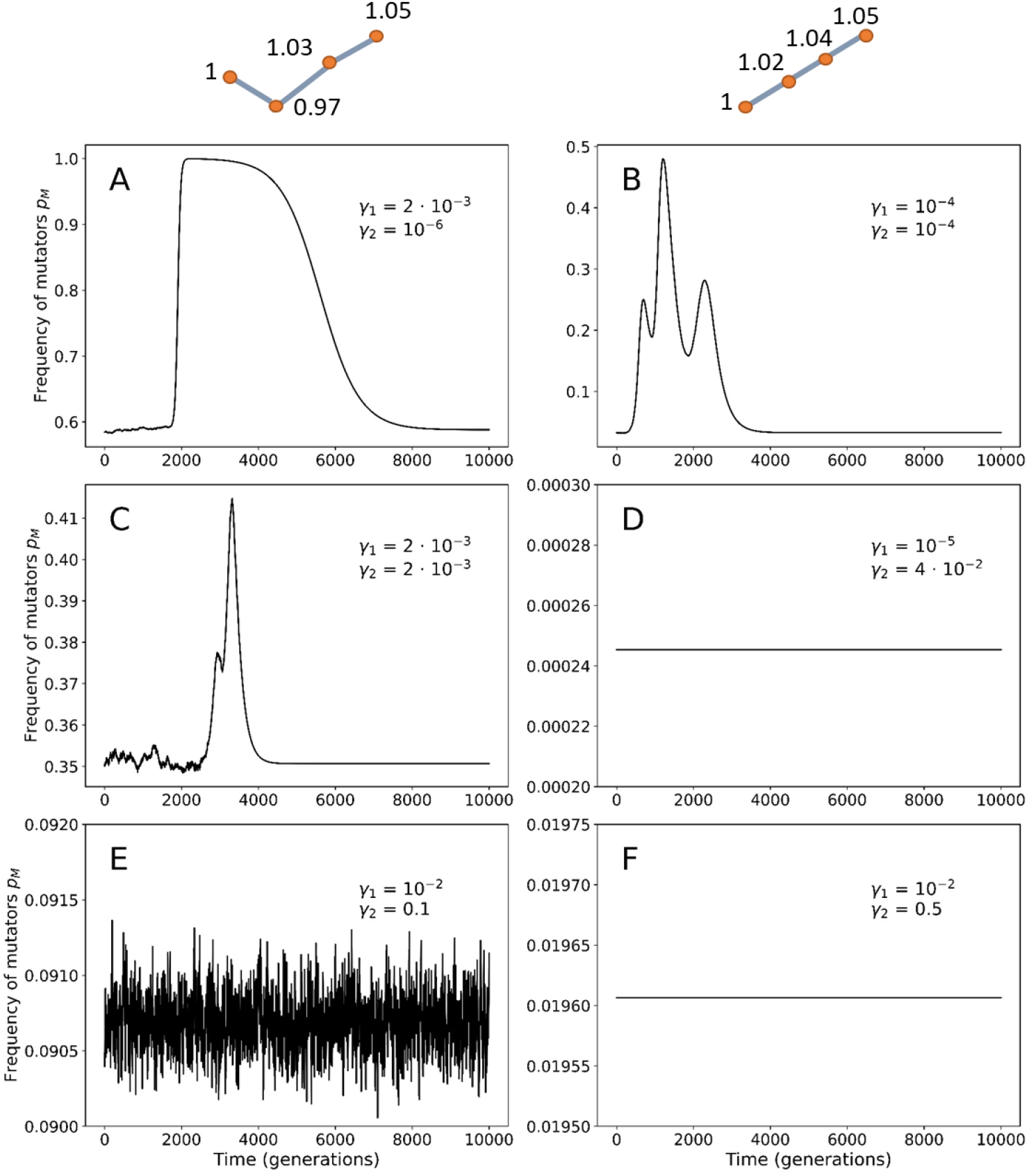
Dynamics of mutator frequency during an adaptive evolution. The frequency of mutators was recorded over 10,000 simulations during adaptation (that is, the appearance and fixation of a fitter genotype). When either of the switching rates is higher than 10^−3^, the mutator frequency barely changes from its mutation-selection balance value. However, when switching rates are lower, the mutator hitchhikes with the adaptive genotype, leading to a transient increase in mutator frequency. For a monotone ascent fitness motifs (right column), each appearance and increase in frequency of an adaptive mutation results in an increase of the mutator frequency, thus facilitating the appearance of the next mutation. We suggest that this phenomenon is responsible for the high adaptation rates observed for *γ*_1_ < *τU*/2 and *γ*_2_ < *τU*/2. Parameters: *U* = 4 · 10^−5^, *τ* = 100, *s* = 0.03, *β* = 5000, *N* = 10^9^.

In **Figure 4**, we can also appreciate that the higher the switching rates, the faster the population returns to mutation-selection balance after the increase in mutator frequency that occurs due to hitchhiking. When either *γ*_1_ > 10^−2^ or *γ*_2_ > 10^−2^, the mutator frequency at mutation-selection balance is undisturbed. However, when *γ*_2_ = 10^−6^, the mutator frequency returns to mutation-selection balance after more than 6,000 generations (defined as being within less than 10^−3^ of the mutation-selection frequency).

If the speed of return of the mutator frequency to the mutation-selection balance is low, the increased frequency of the mutator phenotype associated with genotypes that are intermediate between the wild-type and the adaptive mutant increases the rate of adaptation.

Lastly, we examined whether the fixation time of an adaptive mutant depends on either *γ*_1_ or *γ*_2_. To estimate the fixation time, we measured the number of generations between the appearance of an adaptive genotype and its fixation. We defined fixation exceeding 99% in frequency of the adaptive mutant and measured the time between appearance and fixation for various combinations of the two switching rates. Although some pairs of distributions of times for different switching rates result in a significant p-value, the effect size between is quite small, with Cohen’s *d* less than 0.5 (see **Supplementary Figure 7**).

In summary, stochastic simulations of adaptive evolution have confirmed the analytic results based on mutation-selection balance: the highest adaptation rates are attained for switching rate *γ*_1_ = *τU*/2 when *γ*_2_ < *τU*/2 and *γ*_1_ = *γ*_2_ when *γ*_2_ > *τU*/2.

Moreover, our stochastic simulations have demonstrated an additional mechanism through which non-genetic systems of mutation-rate inheritance can lead to higher rates of adaptation: hitchhiking of the mutator phenotype with the beneficial mutations it generates causes a transient increase in the population-wide mutation rate, facilitating the accumulation of additional beneficial mutations. This phenomenon is observed when the switching rate from mutator to non-mutator is low enough (*γ*_2_ < *τU*/2) and leads higher rates of adaptation than would be expected from the mutation-selection balance analysis, see **Figure 3**.

We have thus studied which combinations of the switching rates from the non-mutator to mutator and vice-versa result in high rates of adaptation. We have identified two mechanisms. The first is a “static” mechanism: some combinations of the parameter space lead to high frequencies of mutator mutants at mutation-selection balance. This situation occurs due to a high frequency of mutator, while maintaining a non-negligible degree of mutator phenotype inheritance that conserves the association between the mutator phenotype and the mutations it generated. We identified this mechanism through the study of mutation–selection balance frequencies of non-mutator and mutator mutants. The second is a “dynamic” mechanism. The hitchhiking of the mutator with the intermediate adaptive mutations facilitates the acquisition of further adaptive mutations. We observed this mechanism through the iteration of our model over what we define as fitness landscape motifs, some of which featured ascending fitness with each subsequent mutation.

## Adaptation on realistic landscapes

In this section, we will determine the relevance of the static and dynamic mechanisms to adaptation on biologically realistic fitness landscapes.

We explore several fitness landscapes: NK landscapes of various ruggedness, which can be used to model adaptive evolution [58], and an empirical landscape derived from the fungus *A. niger* [40], [41]. A description of these landscapes and the choice of the wild-type genotype can be found in the Supplementary material, section “E. Description of complex landscapes” (see **Supplementary Figures 19** and **20**).

We first estimated the mutation-selection balance frequencies of the non-mutator with the initial genotype (*m*_0_), the non-mutator with mutations (*m*_1_), the mutator with the initial genotype (*M*_0_), and the mutator with mutations (*M*_1_) for the NK landscapes (**Supplementary Figures 21**, **22** and **23**), and for the *A. niger* landscape (**Supplementary Figure 24**).

In this section, *m*_0_ is the frequency of non-mutators with the predominant genotype (the “wild-type”), *m*_1_ is the frequency of non-mutators with genotypes other than the predominant genotype (“mutants” of the “wild-type), *M*_0_ is the frequency of mutators with the predominant genotype (the “wild-type”) and *M*_1_ is the frequency of non-mutators with genotypes other than the predominant genotype (“mutants” of the “wild-type). We have *p_S_* = *M*_1_ + *m*_1_ and *p_M_* = *M*_0_ + *M*_1_.

We observe similar patterns to those observed with the simple landscapes (see **Supplementary Figures 2–5**): the frequency of *M*_1_ is highest for *γ*_1_ > *τU*/2 and *γ*_2_ < *τU*/2, and for *γ*_1_ > *γ*_2_ for *γ*_2_ > *τU*/2. We also estimate the frequency of *p_M_* for several parameters sets in **Supplementary Figure 25**, and the association between mutator and mutant phenotypes in **Supplementary Figure 26**. While performing simulations of the stochastic model, we observed comparable patterns to the simple landscape simulations (see **Supplementary Figures 2–5**): a region with high mutator frequency for *γ*_1_ > *τU*/2 and *γ*_2_ < *τU*/2, and *γ*_1_ > *γ*_2_ for *γ*_2_ > *τU*/2, and a negligible frequency of non-mutator single mutants except for *γ*_1_ > 10^−2^ and *γ*_2_ > 10^−2^. These observations indicated to us that the “static” mechanism, that is, the combination of a high frequency of mutators with a high association between the mutator and mutant phenotypes, could be at work also in the case of complex adaptation.

We then simulated the stochastic model on the NK landscapes and the *A. niger* landscape. We plotted the proportion of simulations where more than 99% of the population has the fittest genotype versus the number of generations (**Figure 5** for the *A. niger* landscape and **Supplementary Figure 27–29** for the NK landscapes).

**Figure 5.**
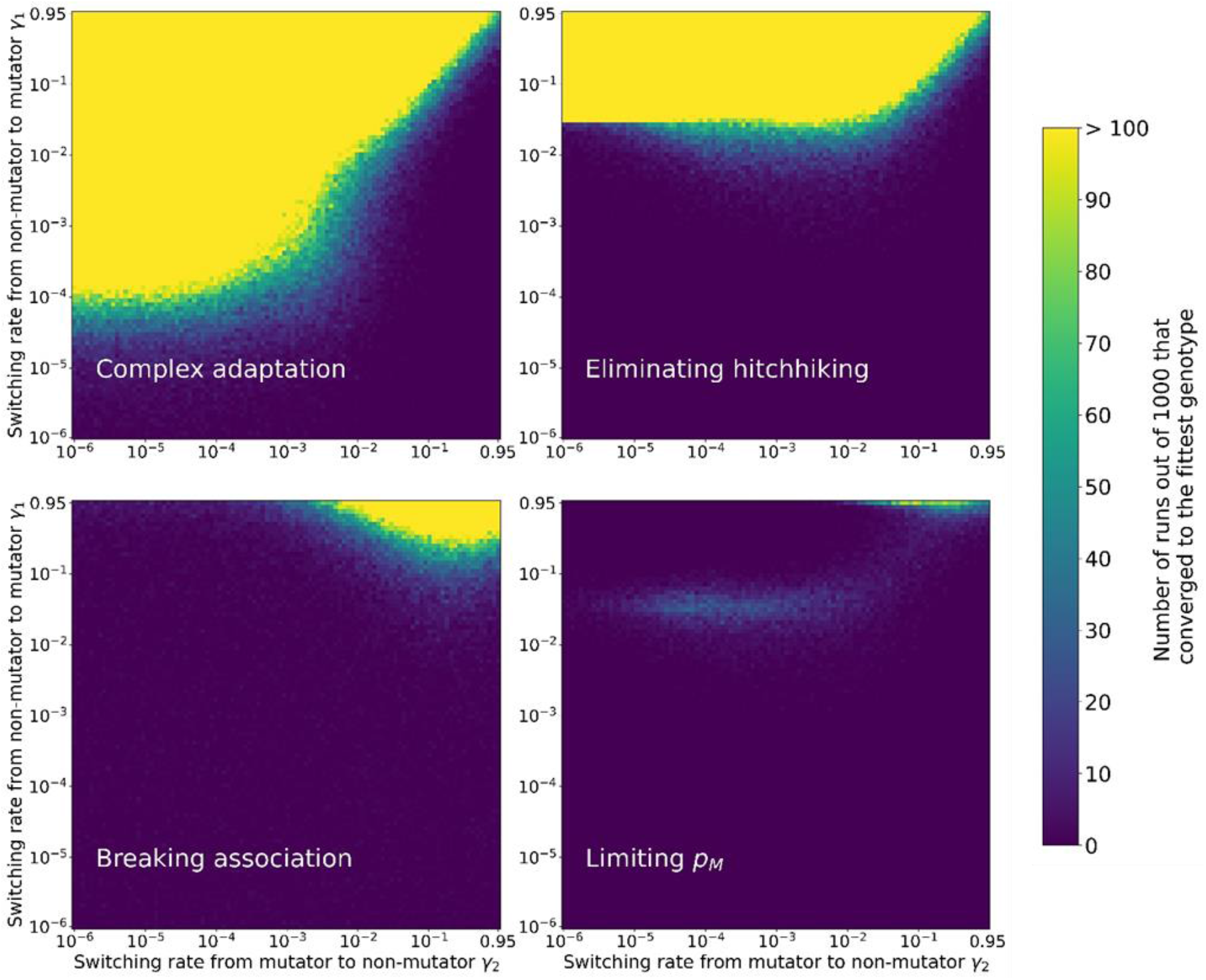
Complex adaptation on *Aspergillus niger* landscape. The proportion of runs out of 1000 that converged upon the fittest genotype in the landscape was recorded after 500 generations. In order to disentangle the different effects of the non-genetic inheritance of the mutation rate on the rate of adaptation, we then rerun the simulation while removing the association of mutator and mutant, limiting the frequency of mutators during the evolution, and eliminating hitchhiking. Note that hitchhiking is also eliminated when the association between mutator and mutant is broken, or the frequency of the mutator limited. We observe that the region with rates of adaptation for *γ*_1_ < *τU*/2 and *γ*_2_ < *τU*/2 disappears when hitchhiking is eliminated (see Supplementary Figure 30 for statistical analysis). Breaking association reduces adaptation except for high *γ*_1_ and *γ*_2_. Limiting the frequency *p_M_* reduces adaptation overall. Parameters: *U* = 4 · 10^−5^, *τ* = 100, *N* = 1000.

We found that the highest rates of adaptation corresponds to the region expected from the frequencies of mutator mutants at mutation-selection balance (**Supplementary Figures 21–25**). We also found an additional region with high adaptation rate for *γ*_1_ < *τU* and *γ*_2_ < *τU* for NK landscapes with *k* = 1 (that is, low ruggedness, see Supplementary section E., Description of realistic landscapes) and *k* = 3 (that is, intermediate ruggedness, see Supplementary section E., Description of realistic landscapes), analogous to the region observed for these parameters for populations adapting over smooth landscapes in **Figure 3**. This is a manifestation of the “standing variation” mechanism. We observe that *γ*_1_ < *τU* and *γ*_2_ < *τU*/2 correspond to high rates of adaptation for NK landscapes with low ruggedness, but not in NK landscapes with high ruggedness. This is a manifestation of the “hitchhiking” mechanism. Hence, we find examples of both of the “standing variation” and “hitchhiking” mechanisms at play in complex adaptation over biologically realistic landscapes.

Next, we determined the relative contribution of the “static” and “dynamic” mechanisms to the rate of adaptation. We simulated the stochastic model while artificially modifying the frequency of the mutator phenotype to eliminate the influence of one of the three conditions that lead to a high adaptation rate: (1) a high mutator frequency – we set the mutator frequency to be half the mutator frequency at mutation-selection balance; (2) high association between the mutator and mutant frequency – we set the association (see Eq. 6) to equal 0 by redistributing mutant genotypes among mutator and non-mutators; and (3) for the dynamic mechanism of the mutator phenotype with an adaptive mutant – we set the mutator frequency to always be the mutator frequency at mutation-selection balance.

### Breaking the association between the mutator and the mutant

High association between mutator and mutant translates into a high frequency of mutator mutants, and in turn increases the rate of adaptation. Hence, we decided to artificially decrease the association between the mutator and the mutant, while keeping the frequency of mutator and mutants, to eliminate this effect from the evolutionary dynamics. At each generation, we artificially break the association (*A* = 0) by setting *m*_0_ = (1 – *p_S_*) · (1 – *p_M_*), *m*_1_ = *p_S_* · (1 – *p_M_*), *M*_0_ = (1 – *p_S_*) · *p_M_* and *M*_1_ = *p_S_* · *p_M_* (see Supplementary Section “H. Complex landscapes: breaking association between mutator and mutant”). The relative frequencies of each mutant within *m*_1_ and *M*_1_ is not modified. Note that when association between the mutator and mutant is broken, hitchhiking is also abolished since the mutator cannot hitchhike on a mutant to increase in frequency.

We observe that the rate of adaptation is significantly reduced compared to the simulation with no changes (**Figure 5** and **Supplementary Figure 30**). The regions with relatively high adaptation rate are obtained for high switching rates *γ*_1_ and *γ*_2_. These are the regions where the frequency of mutators is high. It appears the association between the mutator phenotype and mutant alleles has a strong effect on the adaptation rate. Results for NK landscapes show the same effect (**Supplementary Figure 27–29**).

### Eliminating hitchhiking

To eliminate hitchhiking, we reset the proportion of mutators *p_M_* at each generation to correspond to the value of *p_M_* at mutation-selection balance. Note that in this case, the association between mutator and mutant is also at its mutation-selection balance, that is, it can be different than 0. The relative frequencies of wild-type and mutant genotypes were not modified (see Supplementary Section “G. Complex landscapes: eliminating hitchhiking”).

As described in the previous section, the dynamic mechanism only manifests for both *γ*_1_ < *τU*/2 and *γ*_2_ < *τ*. Indeed, we observed a significantly reduced rate of adaptation for the parameter region *γ*_1_ < *τU* and *γ*_2_ < *τU* (**Figure 5** and **Supplementary Figure 30**). Without hitchhiking, only the parameter region exhibiting high frequencies of mutator mutants at mutation-selection balance showed high rates of adaptation, consistent with the expectation that only the “static” mechanism affects adaptation. On NK landscapes, eliminating hitchhiking reduces the adaptation rate for *γ*_1_ < *τU* and *γ*_2_ < *τU* for low and intermediate ruggedness, but we observe no effect for high ruggedness, further supporting our hypothesis that hitchhiking only occurs on smooth landscapes where successive adaptations occur without the population returning to a mutation-selection balance (**Supplementary Figure 22–24**).

### Reducing the mutator frequency

In the previous section, we observed that high adaptation rates are associated with high mutator frequency, which in turn is associated with high mutator mutant frequency. To reduce the mutator frequency, we reset the proportion of mutators *p_M_* to half of its mutation-selection balance value at each generation. The relative frequencies of wild-type and mutant genotypes were not modified (see Supplementary Section “F. Complex landscapes: reducing the mutator frequency”). Note that in this simulation, hitchhiking is also abolished since the mutator cannot hitchhike on a mutant to increase in frequency. The rate of adaptation is reduced, mostly for the areas where *p_M_* > 0.5 at mutation-selection balance (**Figure 5** and **Supplementary Figure 30**). However, we still observe relatively high adaptation rates with *γ*_1_ = *τU* when *γ*_2_ < *τU* and *γ*_1_ = *γ*_2_ when *γ*_2_ > *τU*. Results of simulation on NK landscapes are similar (**Supplementary Figure 27–29**).

In summary, we studied how the “static” and “dynamic” mechanisms intervene in the evolution over realistic landscapes. The “static” mechanism refers to the advantage in adaptation stemming from the frequency of mutator mutants at mutation-selection balance. The frequency of mutator mutants is high when the frequency of mutators is high and when the frequency of mutants that also mutators is high compared to their expected frequency given the frequency of mutants and mutators. Removing either of them results in lower adaptation when *γ*_2_ < *τU* and *γ*_1_ > *τU* and when *γ*_2_ > *τU* and *γ*_1_ > *γ*_2_. The “dynamic” mechanism manifests on landscapes with low ruggedness and *γ*_1_, *γ*_2_ < *τU*. When removing the effect of hitchhiking, we observe low adaptation for specifically *γ*_1_, *γ*_2_ < *τU*.

## Discussion

### The potential advantage of non-genetic inheritance of the mutation rate

We have shown that the combination of the switching rates that lead to high adaptation rates on simple landscapes also result in high adaptation rates on complex, more biologically realistic landscapes. The result is maintained also when considering gradual transition between the non-mutator and mutator phenotypes (see **Supplementary Figure 1, 10 and 11**). The advantage combination of switching rates corresponding to non-genetic inheritance of the mutation rate in the rate of adaptation seems to come from two mechanisms, a static mechanism relying on a high frequency of mutator mutants that exists even at mutation-selection balance, and a dynamic mechanism that manifests itself while hill climbing of fitness landscape.

The static mechanism leads to high rates of adaptation for *γ*_1_ = *τU*/2 when *γ*_2_ < *τU*/2 and *γ*_1_ = *γ*_2_ when *γ*_2_ > *τU*/2. For highest rates of adaptation to be achieved, the inheritance of the mutation rate phenotype needs to be non-negligible, hence *γ*_1_ < 10^−2^ and *γ*_2_ < 10^−2^. For smooth landscapes with potential hill climbing, the dynamic mechanism also leads to high adaptation rates for *γ*_2_ < *τU*/2 and 10^−2^ < *γ*_1_ < *τU*/2. These values and mechanisms are summarized in **Figure 6**.

**Figure 6.**
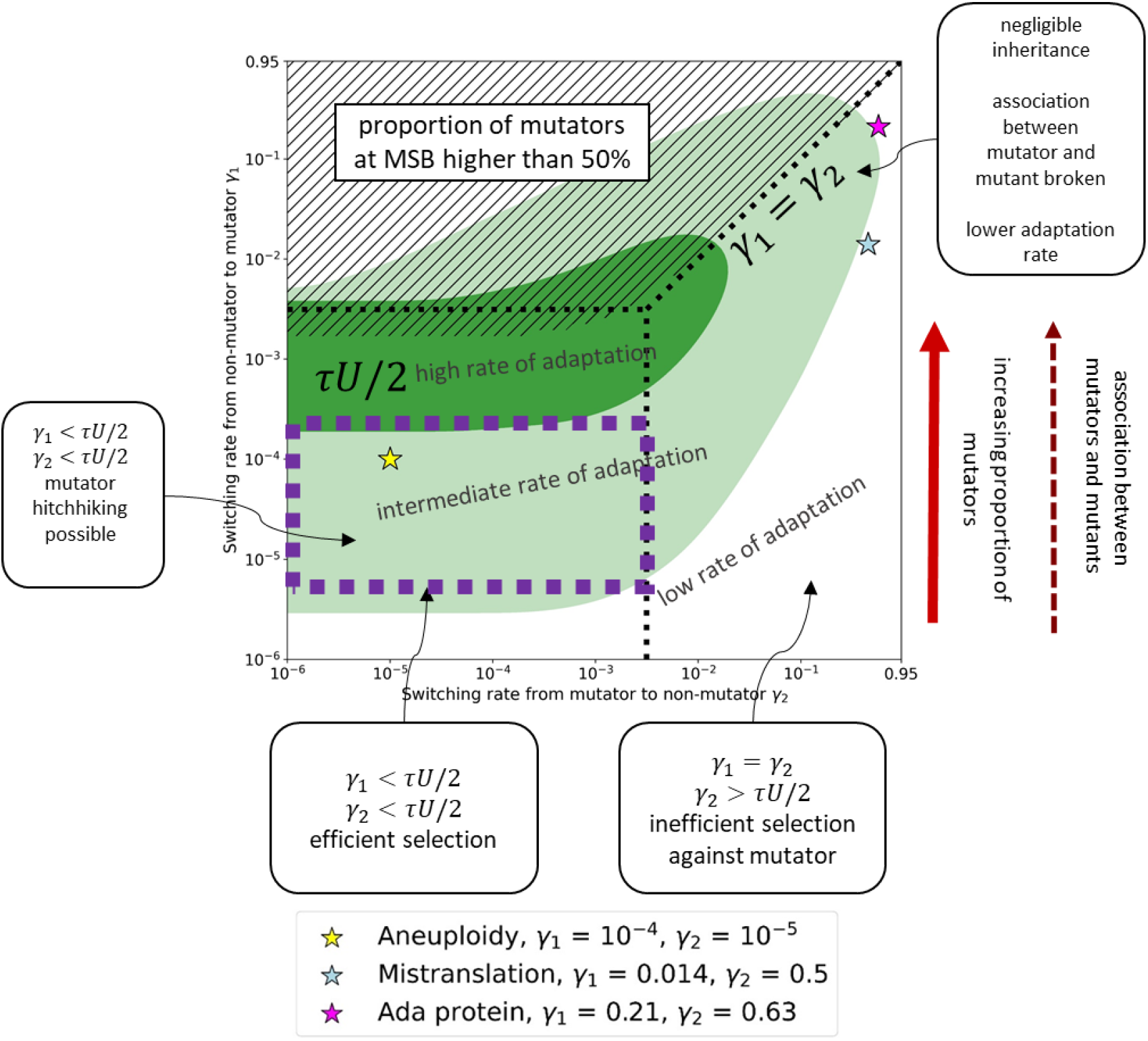
Graphical summary of the rate of adaptation under non-genetic inheritance of the mutation rate. As the switching rate from non-mutator to non-mutator increases, the frequency of mutators at mutation-selection balance increases as well. This creates an emergent association between mutator and mutant phenotypes that is higher than expected by chance. When *γ*_2_ < *τU*/2, the region with highest adaptation rate is around *γ*_1_ = *τU*/2. When *γ*_2_ > *τU*/2, the region with highest adaptation rate is on the diagonal, corresponding to *γ*_1_ = *γ*_2_, except for when mutator phenotype inheritance is very weak, i.e., *γ*_1_ > 0.01 and *γ*_2_ > 0.01. in which case adaptation rate is not as high. The hashed region corresponds to switching rate combinations that result in mutator frequency above 50%, hence excluded from the model. The three coloured stars represent the three systems of non-genetic inheritance of the mutation rate: aneuploidy, cytoplasmic inheritance of mistranslated proteins, and the Ada system from *E. coli*. All three systems are expected to produce high adaptation rates.

After performing simulations attempting to eliminate each of these mechanisms to assess their relative contributions in complex adaptation we deduce that the static mechanism seems to be decisive, since abolishing the association between mutator and mutants drastically decreases the rate of adaptation. The dynamic mechanism is only important for a subset of the parameter space, for *γ*_1_ < *τU*/2 and *γ*_2_ < *τU*/2, and for smooth landscapes.

### Evolution of intermediate switching rates

Although some combinations of switching rates maximize the adaptation rate, it is not clear that they can be selected for when competing against other combinations of switching rates. First, the proportion of mutants for the adaptation-optimal pairs of switching rates is high, higher than for subpopulations with adaptation-suboptimal combinations of switching rates (see **Supplementary Figures 2–5**). Second, the advantage of a subpopulation with an adaptation-optimal switching rate depends on the mutation supply. The adaptation rate of a subpopulation of size *N*_1_ with a switching rate *γ*_1_ resulting in an appearance rate *q*_1_ will be higher than the adaptation rate in a subpopulation of size *N*_2_ with a given switching rate *γ*_2_ resulting in an appearance rate *q*_2_ if 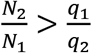 (Eq. 5). Therefore, selection for an allele that induces an intermediate switching rates will depend on its frequency in the population, similar to the case of mutator alleles [59].

### Mutation rate inheritance and phenotype switching

Switching between a mutator and a non-mutator phenotype is a type of phenotype switching [60]. The increased adaptability of populations with adaptation-optimal pairs of the two switching rates, compared to other pairs, is in part due to the high rate of mutator generation, which balances the strength of selection against mutators at the mutation-selection balance. Generation of a phenotype that is disadvantageous in the short term but potentially advantageous in the long term is known as bet-hedging [60] and has been implicated as driving the evolution of phenotype switching mechanisms [60]–[63].

We suggest that the mechanisms we identified to be implicated in high rates of adaptation for some pairs of switching rates could be at play also in other systems in which a specific phenotype has a short-term disadvantage but long-term advantage. High switching rate from wild-type to phenotype counters negative selection, and high association between the phenotype allows for the manifestation of the long-term advantage. In such cases, if the same phenotype may also be advantageous if it occurs in consecutive generations of the same lineage, and therefore a not too high switching rate (in our model, less than 10^−2^) is beneficial in maintaining a correlation between the phenotype of parent and offspring.

### Experimental future directions

Our results show theoretically that non-genetic inheritance of the mutation rate could result in higher rates of adaptation as a general rule. Interestingly, current methods for estimating the mutation rate, such as mutation accumulation [64] and fluctuation assays [45], require multiple generations along which mutation rate is measured. Hence, they implicitly assume that the mutation rate is constant across generations and individuals. This is why diversity and non-genetic inheritance of the mutation rate has not been observed until recently [11], [65], [66]. Our study provide an evolutionary rationale for these observed mutation rate dynamics and it calls for a new focus on non-genetic factors influencing the mutation rate, and its potential impact on adaptative evolution.

**Table 2.** Estimates of switching rates *γ*_1_ and *γ*_2_ for three considered mechanisms for non-genetic inheritance of the mutation rate.

| Switching rate | Estimate | References |
| --- | --- | --- |
| Ada protein |  |  |
| Non-mutator to mutator $\gamma_1$ | 0.21 | [11] |
| Mutator to non-mutator $\gamma_2$ | 0.63 | |
| Aneuploidy |  |  |
| Non-mutator to mutator $\gamma_1$ | $10^{-4}$ | [27], [50]–[53] |
| Mutator to non-mutator $\gamma_2$ | $10^{-5}$ | |
| Cytoplasmic inheritance of mistranslated proteins |  |  |
| Non-mutator to mutator $\gamma_1$ | 0.014 | [10], [14], [55], [56], [77] |
| Mutator to non-mutator $\gamma_2$ | 0.5 | |

## Data availability statement

The code necessary to reproduce the results and the plots presented in this paper is available at https://github.com/gabriela3001/revision_phenotypic_mrate, with Zenodo id: https://doi.org/10.5281/zenodo.7584835.

## Acknowledgments

YP holds the Ben May Professorial Chair, and is a Kimmel Investigator.

YR acknowledges the Minerva Foundation and the Israel Science Foundation (552/19) for grant support.

We thank Eytan Domany, Ariel Amir, Naama Barkai, and Tal Simon, for discussions and comments.

## Appendix

### A. Simple landscape

The mutation transition probability is *u_j→g_*, where *j* and *g* are genotypes, determines the effect of mutation on the genotype probabilities (Eq. 1). The construction of the mutation transition probability is described in the main text. Here, we provide a formal description. Note that we assume no back mutations occur. In the following we use the mutator phenotype mutation rate *τU;* for the non-mutator, we set *τ=1*. The per-locus mutation rate is *μ* = *U/L*. The number of deleterious mutations in the background loci is *k* = 0,1,2,….

Thus, the probability to transition from genotype *j* to genotype *g*, *u_j→g_*, is described by **Table A1**, where the source genotype *j* is given in the row, and the target genotype *g* is given in the column. We do not distinguish between *Ab* and *aB* as we assume they have equal fitness values.

**Table A1.** Probabilities to transition from genotype *j* to genotype *g*, *u_j→g_*.

| $u_{j \rightarrow g}$ | $g = ab/k+l$ | $g = Ab/k+l$ or $aB/k+l$ | $g = AB/k+l$ |
| --- | --- | --- | --- |
| $j = ab/k$ | $(1 - \tau\mu)^2 \cdot \frac{e^{\tau U} (\tau U)^l}{l!}$ | $2\tau\mu(1 - \tau\mu) \frac{e^{\tau U} (\tau U)^l}{l!}$ | $(\tau\mu)^2 \frac{e^{\tau U} (\tau U)^l}{l!}$ |
| $j = Ab/k$ or $aB/k$ | 0 | $(1 - \tau\mu) \cdot \frac{e^{\tau U} (\tau U)^l}{l!}$ | $\tau\mu \cdot \frac{e^{\tau U} (\tau U)^l}{l!}$ |
| $j = AB/k$ | 0 | 0 | $\frac{e^{\tau U} (\tau U)^l}{l!}$ |

### B. Calculating the switching rate from mutator to non-mutator *γ*_2_ for the *Ada* protein

We define a Markov chain with two states: cells with zero Ada molecules denoted by *x* and cells with non-zero Ada molecules denoted by *y*.

The transition matrix *T* is given by

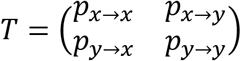

where *p_x→x_* = 1 – *p_x→y_* and *p_y→y_* = 1 – *p_y→x_*. In [11], the probability of transitioning from zero to one (or more) Ada molecules is fitted to a Poisson distribution with average equal to 1. Hence, we have *p_x→x_* = 0.37 and *p_x→y_* = 1 – *p_x→x_* = 0.63.

The stationary distribution of this Markov chain is denoted by (*x*\*, *y*\*) with *y*\* = 1 – *x*\*. Empirical evidence [11] suggests that (*x*\*, *y*\*) ≈ (0.25,0.75). To estimate *p_y→y_*, we solve

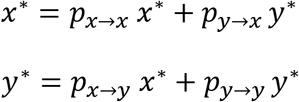

to obtain

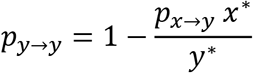

Hence, the complete transition matrix is

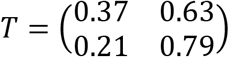

### C. Bounds on the population size

We consider *N*, the population size, to be large enough so that individuals with a single mutation in the major loci are present, but small enough so that individuals with two mutations in the major loci (hence already adapted to the new environment) are absent.

Let us first consider the first condition. The lowest frequency of mutants is achieved when the whole population is non-mutator. Therefore, the upper bound for the first condition is set by considering a case where all individuals have the non-mutator phenotype. According to [67], the frequency of wild type individuals at mutation-selection balance is *e^−U/S^*. We then multiply by the mutation-selection balance of single mutants at the major loci, 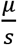. Thus, we find that the expected number of single mutant individuals without deleterious mutations in the background loci is 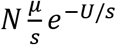. Setting this to be less than one and rearranging, we obtain 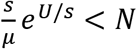.

Now let us consider the second condition. The highest frequency of double mutants would be achieved when the whole population is mutator. Therefore, the lower bound for the second condition is set by considering a case where all individuals have the mutator phenotype. By the same argument as for the first condition, we find 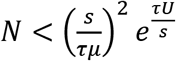 with 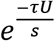 the proportion of wildtype individuals in an all-mutator population and 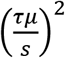 the frequency of double mutants in the major loci, both at a mutation selection balance.

Combining these conditions, we have

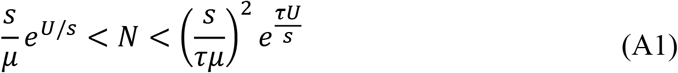

### D. Fixation of an adaptive genotype

According to Eshel [68], in a large population with weak selection, the fixation probability *ρ_F_* of a double mutant once it appears in a single copy depends only on the selection coefficient *s* and the double mutant advantage *H*, namely

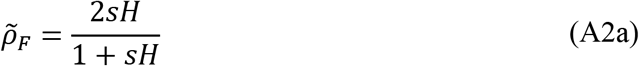

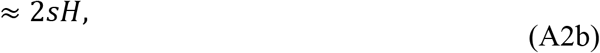

where Eq. A2b applies when *sH* is much smaller than 1.

**Supplementary Figure 6** shows a comparison of the two approximations in eq. A2 to results from stochastic simulations obtained by counting the number of fixation events after the appearance of an adaptive genotype.

We notice that Eq. A16b is independent from the switching rate *γ*. Hence, the adaptation rate will be proportional to the appearance rate (see **Supplementary Figure 6**). We therefore consider the appearance rate as a proxy for the adaptation rate in the main text.

### E. Description of complex landscapes

#### NK landscapes

The NK landscape is commonly used in the study of epistatic interactions [39]. It has two parameters: *n* for the number of bi-allelic loci in the genotype and *k* for the number of loci each locus interacts with. The main advantage of the NK landscape is its biological interpretation and its tunable ruggedness [69]. Here, the genotype consists of *n*=6 bi-allelic loci, which results in 2^6^ = 64 genotypes.

We chose *n* = 6 in order to allow for multidimensionality, while maintaining reasonable computation times.

To construct an NK landscape, we first generate all possible *k*-bit strings (bit strings of length *k*) and assign to each of them a random fitness value between 0 and 1, sampled from a continuous uniform distribution. Genotype *g* is a *n*-bit string, and its fitness *w_g_* is the sum of the fitness effects of the *k*-bit strings that *g* contains. A locus thus influences the genotype fitness according to the number of *k*-bit strings that contain it, which increases with *k*.

Hence, the ruggedness of the NK landscape increases with *k* [40]. We consider three NK landscapes with low, intermediate, and high ruggedness corresponding respectively to *k* = 1, *k* = 3, and *k* = 5. See **Figs. X** and **X** and **Table A2** for properties of the constructed landscapes.

#### Empirical fitness landscape from *Aspergillus niger*

We examine the insights gained on two-peak and NK landscapes with an empirical fitness landscape [39] measured with mutants in 8 genes of *Aspergillus niger* [40]. We chose this landscape because: (i) it is complete with fitness measurements of all possible 256 genotype combinations, so we can avoid the interpolation of fitness values of missing genotypes; and (ii) fitness was measured as growth rate relative to the wild type, as opposed to other studies that quantify some proxy phenotype for fitness such as fluorescence or DNA binding [16], [70], [71].

The genotype consists of eight bi-allelic loci, each with either the wild type or the mutant allele. de Visser *et al*. [40] engineered 256 genotypes to bear all possible combinations of wild type/mutant alleles in these eight loci. The loci are: *fwnA1* (fawn-colored conidiospores); five auxotrophic markers, *argH12* (arginine deficiency), *pyrA5* (pyrimidine deficiency), *leuA1* (leucine deficiency), *pheA1* (phenyl-alanine deficiency), and *lysD25* (lysine deficiency); and two resistances, *oliC2* (oligomycin resistance) and *crnB12* (chlorate resistance).

de Visser *et al*. [40] estimated fitness by measuring the growth rate of each strain relative to the growth rate of the wild type strain (i.e., with eight wild type alleles). Out of 256 genotypes, 70 genotypes were lethal (fitness is zero). The landscape is rugged: it contains 15 local maxima (including the global maximum). See **X** and **X** for properties of the landscape.

### F. Complex landscapes: eliminating hitchhiking

The stochastic was run following the usual mutation – phenotypic switching – selection – drift scheme. Right before the drift step, the population vector was recalculated as follows:

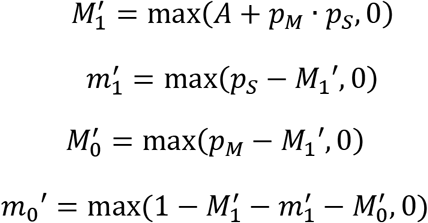

The genotype 0 is the most abundant genotype at that generation. All the mutants’ (mutants are all genotypes that are not the genotype 0) frequencies are kept identical relative to each other, but their sum is set to 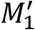 (for mutator mutants) or 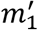 (for non-mutator mutants).

### G. Complex landscapes: limiting

The stochastic was run following the usual mutation – phenotypic switching – selection – drift scheme. The desired frequency of mutators is equal to *M*. Right before the drift step, the population vector was recalculated as follows:

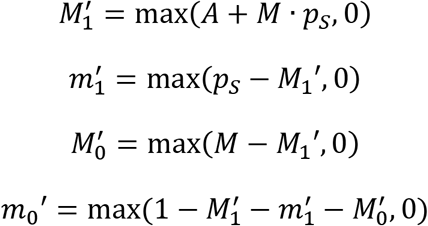

The genotype 0 is the most abundant genotype at that generation. All the mutants’ (mutants are all genotypes that are not the genotype 0) frequencies are kept identical relative to each other, but their sum is set to 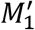 (for mutator mutants) or 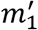 (for non-mutator mutants).

### H. Complex landscapes: breaking the association between mutators and mutants

The stochastic was run following the usual mutation – phenotypic switching – selection – drift scheme. Right before the drift step, the population vector was recalculated as follows:

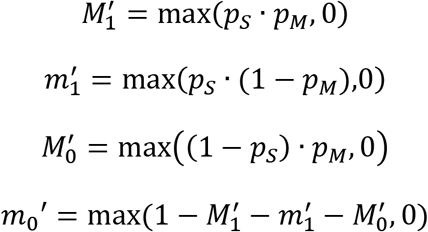

The genotype 0 is the most abundant genotype at that generation. All the mutants’ (mutants are all genotypes that are not the genotype 0) frequencies are kept identical relative to each other, but their sum is set to 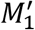 (for mutator mutants) or 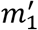 (for non-mutator mutants).

## Figures

**Supplementary Figure 1.**
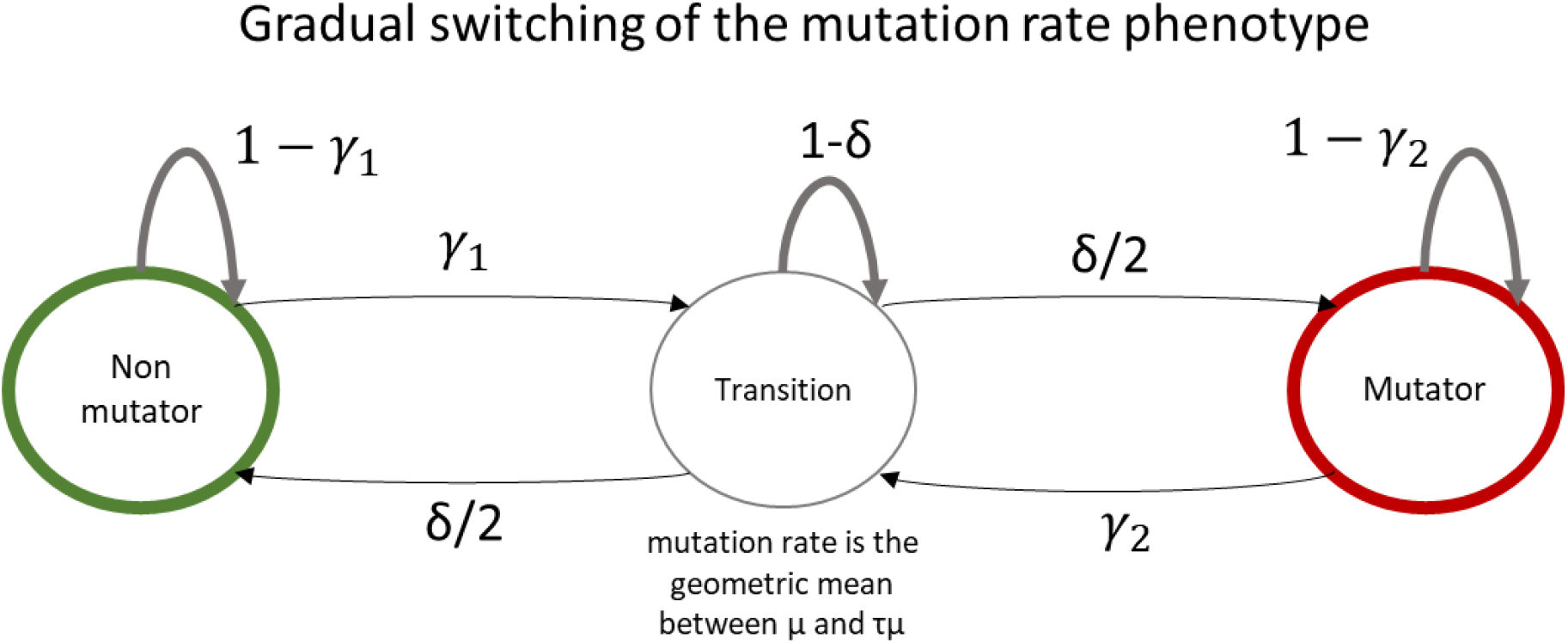
Model extension for gradual switching of the mutation rate phenotype. In this model extension, we introduce an additional, transitional genotype with mutation rate equal to the geometric mean between the non-mutator and the mutator mutation rate. The switching rate *γ*_1_ indicates the switching from the non-mutator to this transition phenotype; a new parameter, *δ*/2, governs the switching from the transition to the mutator phenotype. By symmetry, we also have the switching rate *γ*_2_ from the mutator to the transition phenotype and *δ*/2 the probability of switching from the transition phenotype to the non-mutator phenotype.

**Supplementary Figure 2.**
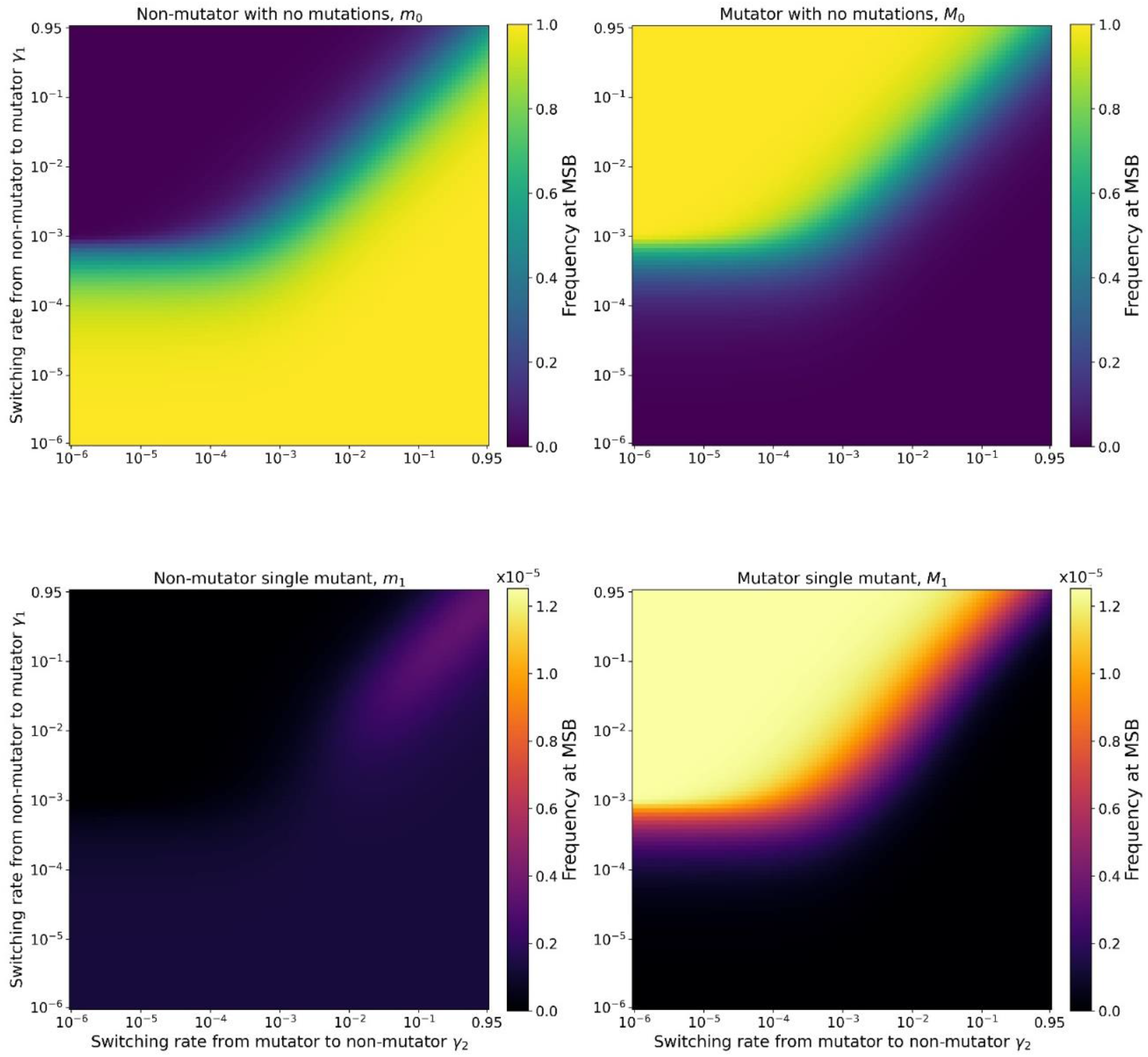
Proportion of non-mutator with no mutations, mutator with no mutations, non-mutator single mutant, and mutator single mutant at mutation-selection balance. As expected, the proportion of single mutants is several orders of magnitude lower than the proportion of mutators. The frequencies of mutator with no mutations and mutator single mutant are in correspondence, with highest frequencies observed for high switching from non-mutator to mutator. The frequencies of non-mutator single mutant are in general very low, except when the inheritance of the mutation rate is negligible (*γ*_1_ > 0.01 and *γ*_2_ > 0.01) and *γ*_1_ = *γ*_2_. Parameters: *U* = 4 · 10^−5^, *τ* = 100, *s* = 0.03, *β* = 5000.

**Supplementary Figure 3.**
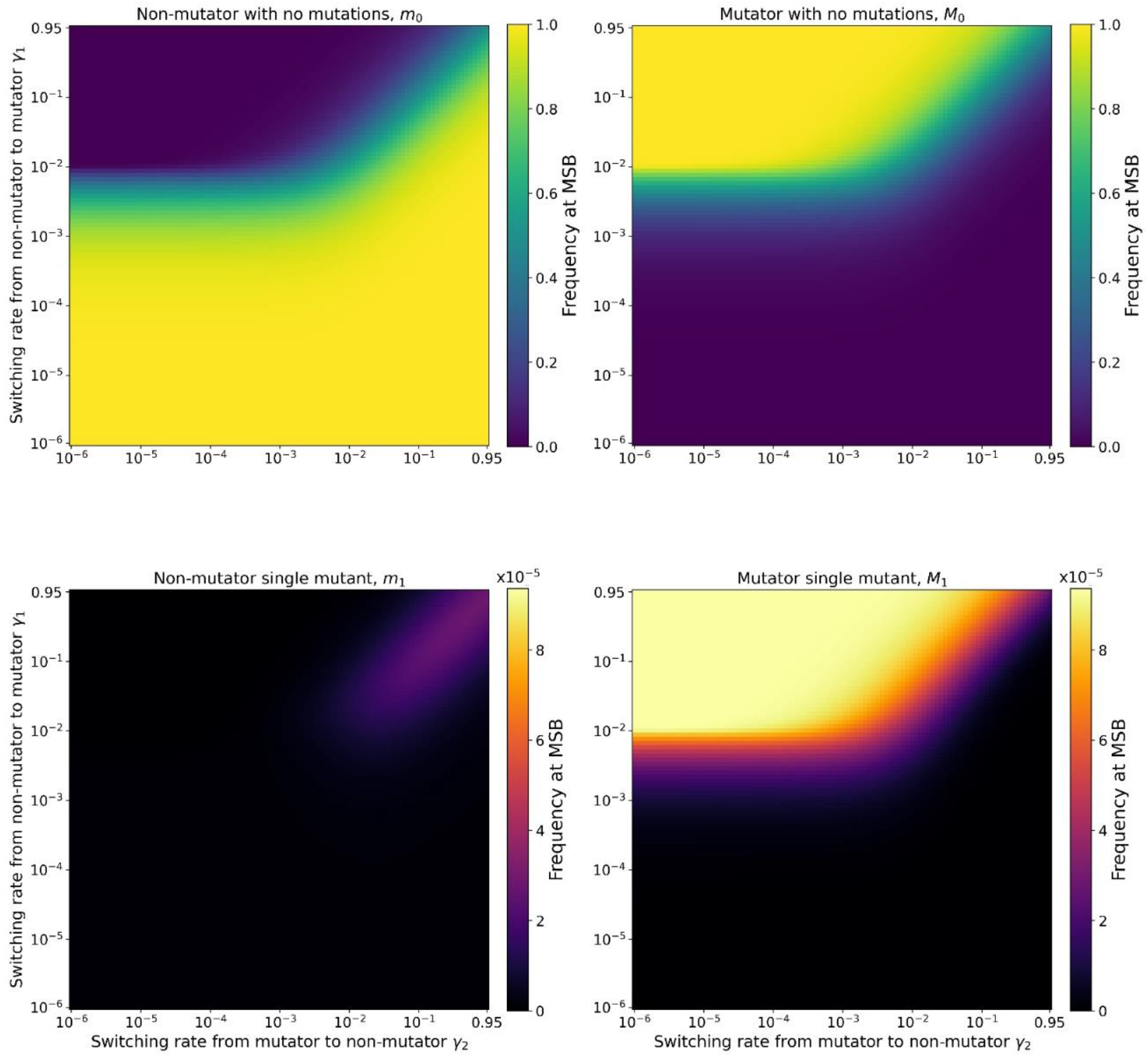
Proportion of non-mutator with no mutations, mutator with no mutations, non-mutator single mutant, and mutator single mutant at mutation-selection balance. Same as Supplementary Figure 2, but with *τ* = 10.

**Supplementary Figure 4.**
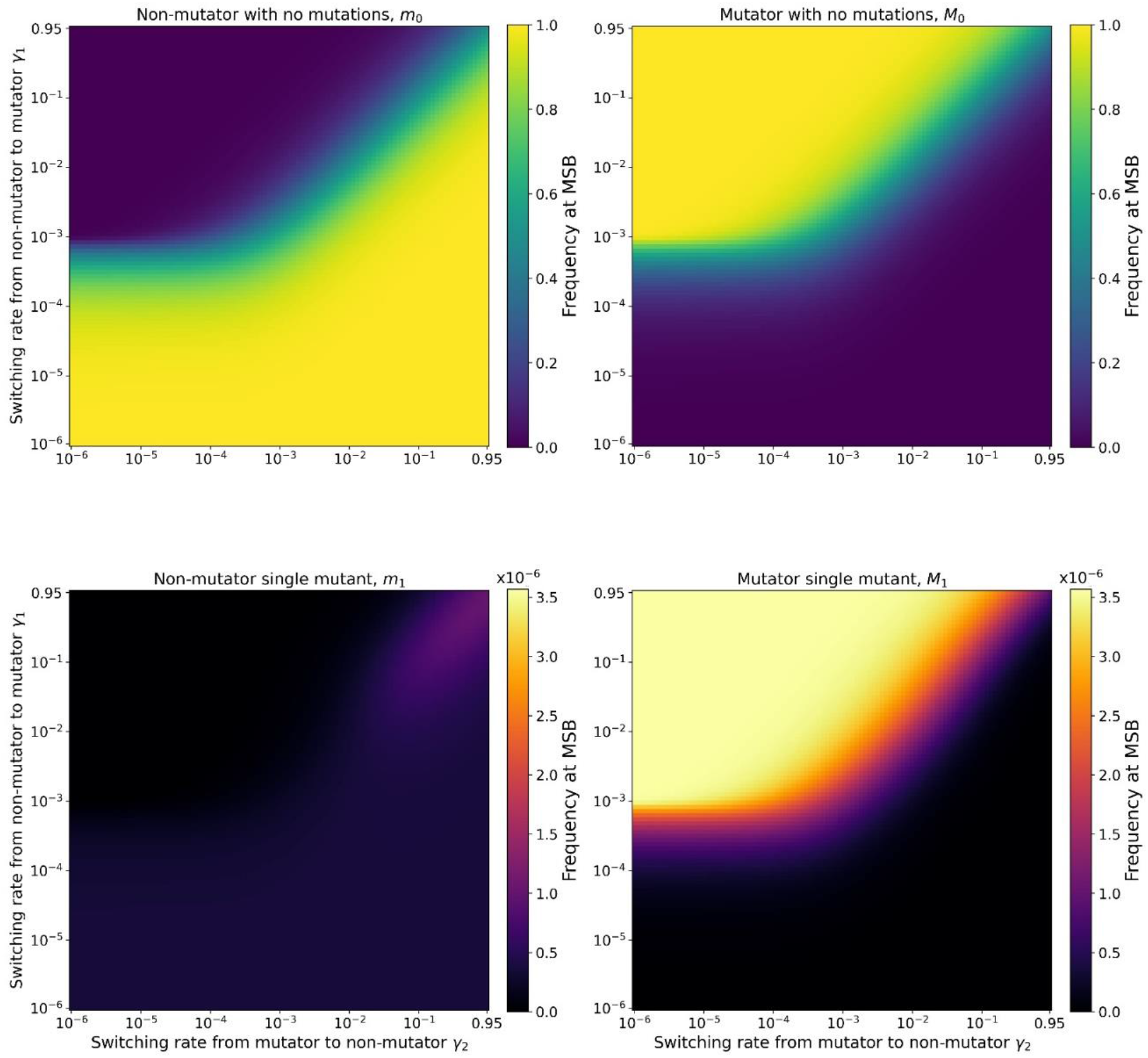
Proportion of non-mutator with no mutations, mutator with no mutations, non-mutator single mutant, and mutator single mutant at mutation-selection balance. Same as Supplementary Figure 3, but with *s* = 0.1.

**Supplementary Figure 5.**
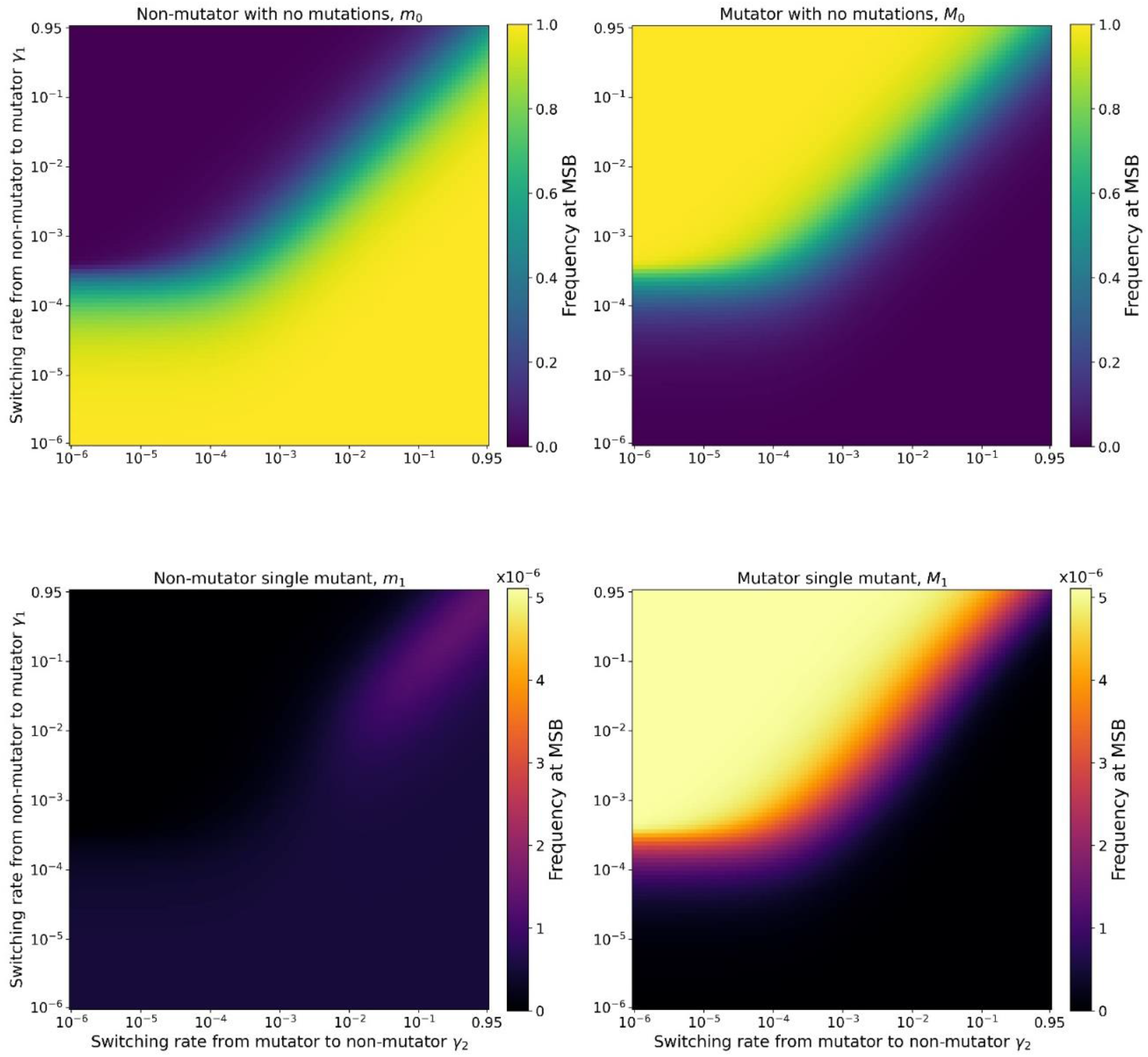
Proportion of non-mutator with no mutations, mutator with no mutations, non-mutator single mutant, and mutator single mutant at mutation-selection balance. Same as Supplementary Figure 3, but with *U* = 10^−4^.

**Supplementary Figure 6.**
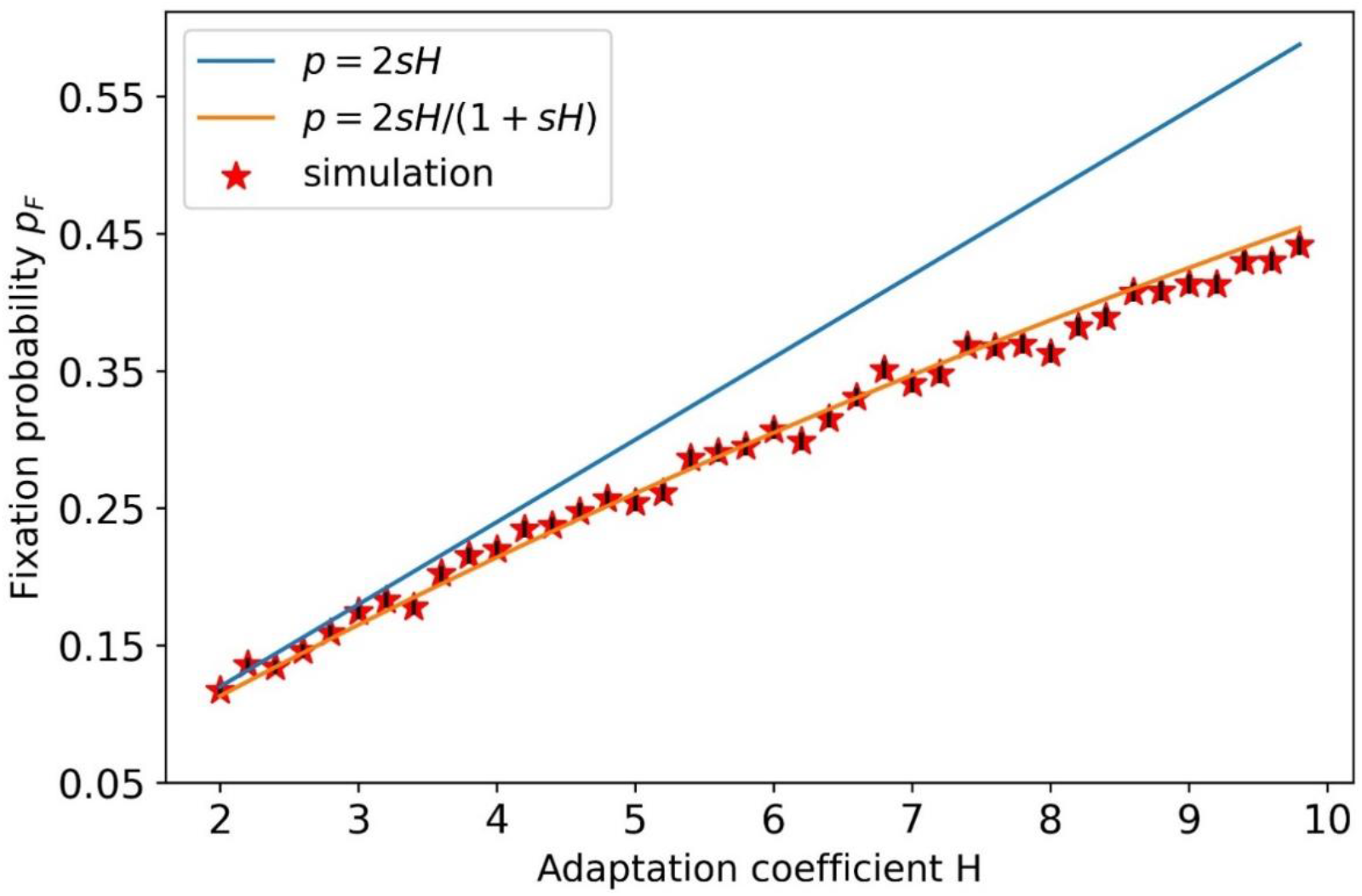
Fixation probability of a rare beneficial genotype. Red stars represent the frequency of fixations in n = 5000 events of adaptive genotype appearance. Error bars (too small to see) show the estimated error 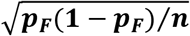. The analytic approximation in Eq. A2a (orange) explains ***R*^2^ = 0.9968** of the variance in the simulation results, while the approximation Eq. A2b (blue) explains ***R*^2^ = 0.9919**. Here, s = 0.03.

**Supplementary Figure 7.**
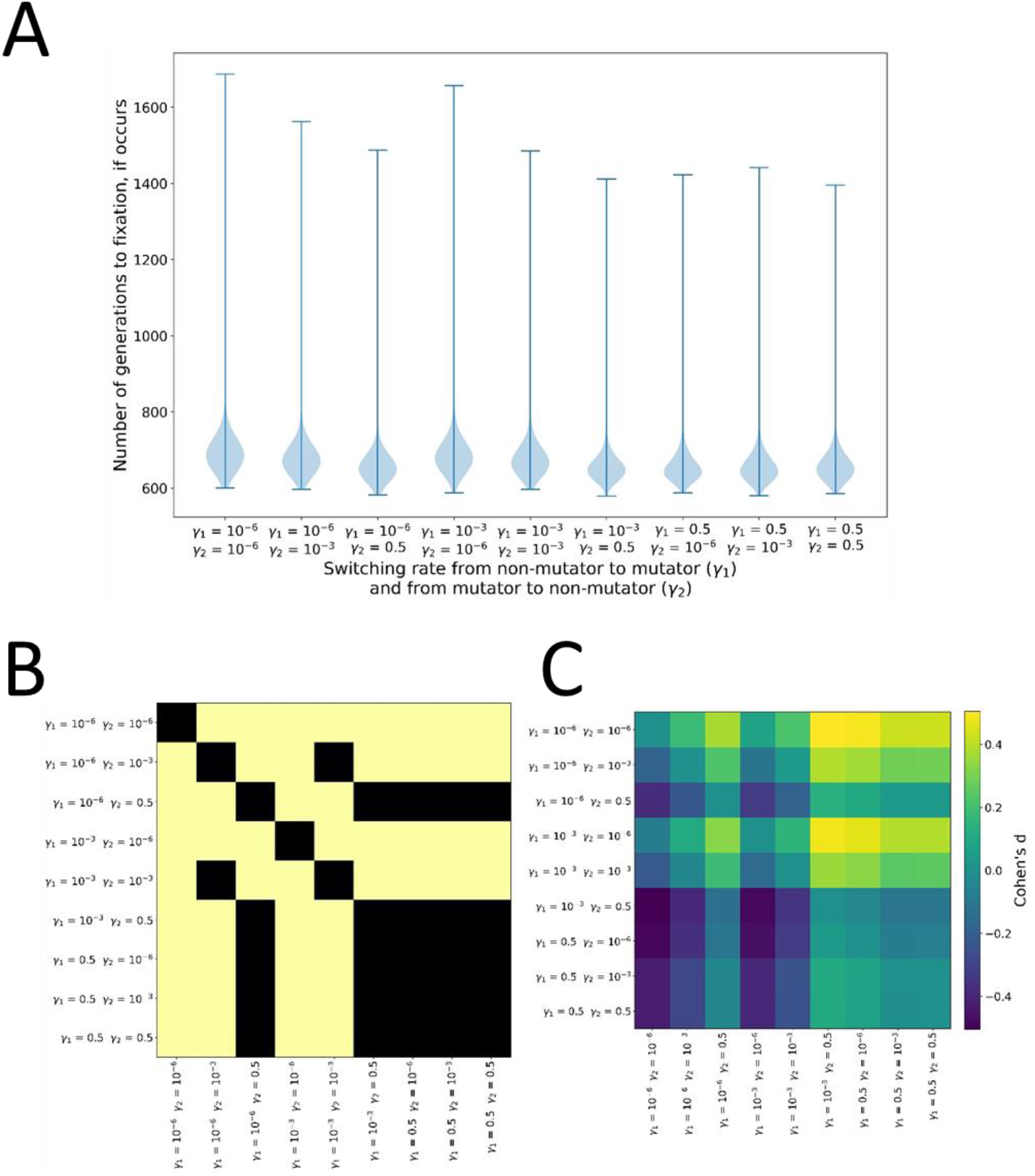
**(A) Time to fixation for pairs of *γ*_1_ and *γ*_2_.** The time to fixation follows a right-tailed distribution that is not strongly dependent on the values of *γ*_1_ and *γ*_2_. The adaptive mutant had a 6% advantage over the wild-type. **(B) Significance of all pairwise comparisons of time distributions.** A Mann-Whitney test was performed with significance threshold set at 0.01, and a Bonferonni correction was applied to correct for multiple comparisons. Distributions where one switching rate equals 0.5 seem to not be significantly different from one another. Yellow represents statistical significance, black represents lack of statistical significance. **(C) Effect size of the difference between the pairs of the time distributions**. Calculated with Cohen’s *d*. Maximal effect size is less than 0.5, which is quite small.

**Supplementary Figure 8.**
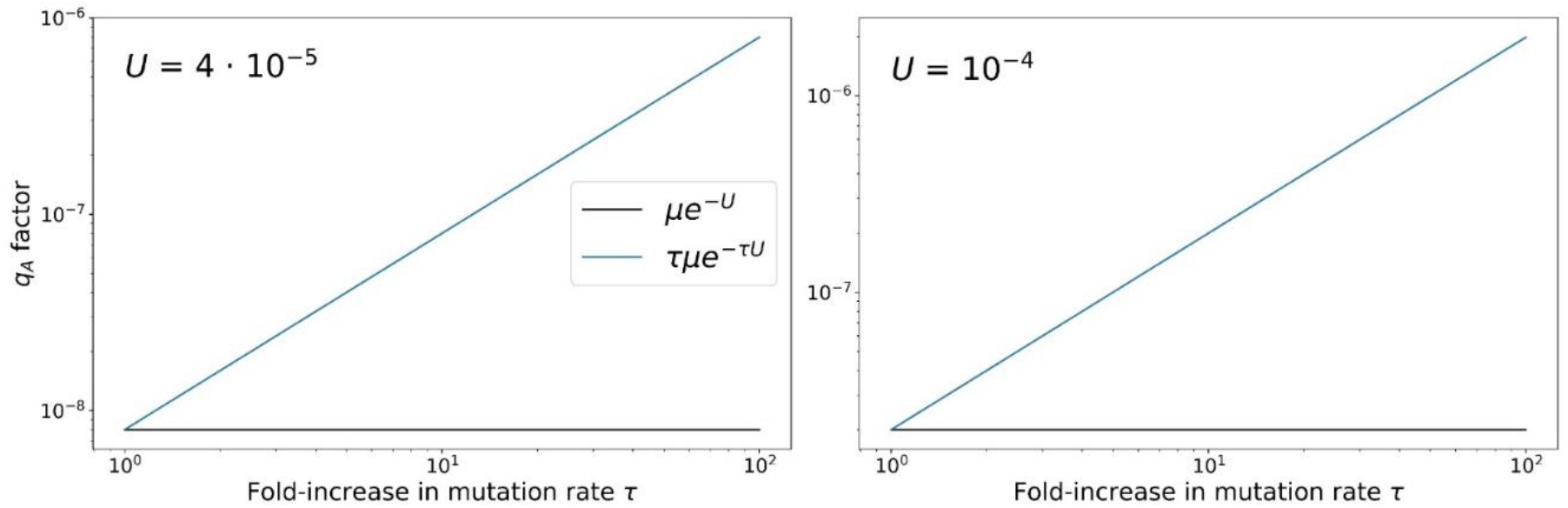
Comparison of *μ e^−U^* and *τμ e^−U^* versus *τ*. We notice that *μ e^−U^* is negligible with regards to *τμ e^−τU^* for almost all of the considered values of τ, the fold-increase in mutator mutation rate.

**Supplementary Figure 9.**
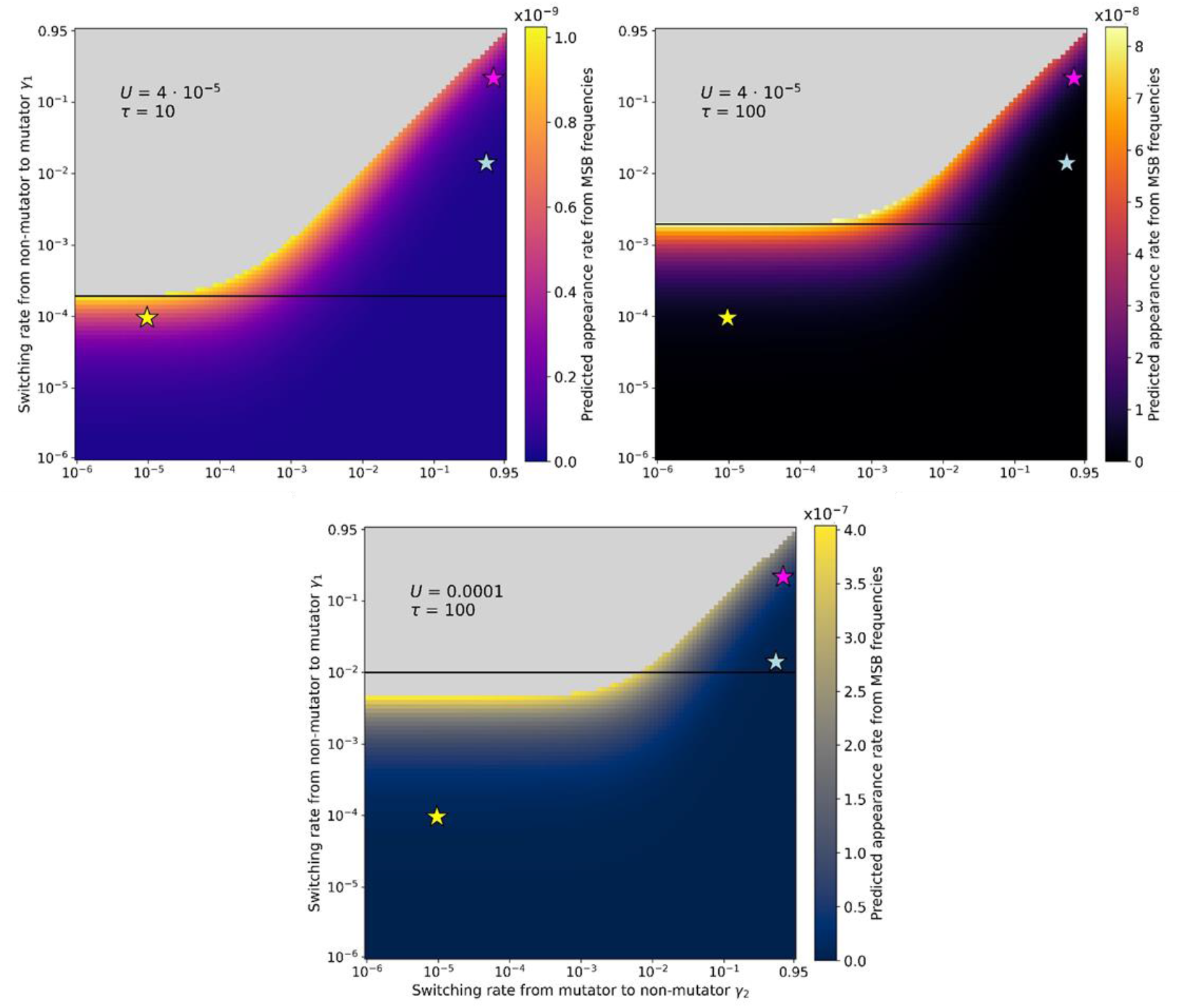
Predicted rate of adaptation based on mutation-selection balance frequencies of mutators and mutants. Same as Figure 2, but for additional parameter sets of *U* and *τ*. Parameters: *s* = 0.03, *β* = 5000.

**Supplementary Figure 10.**
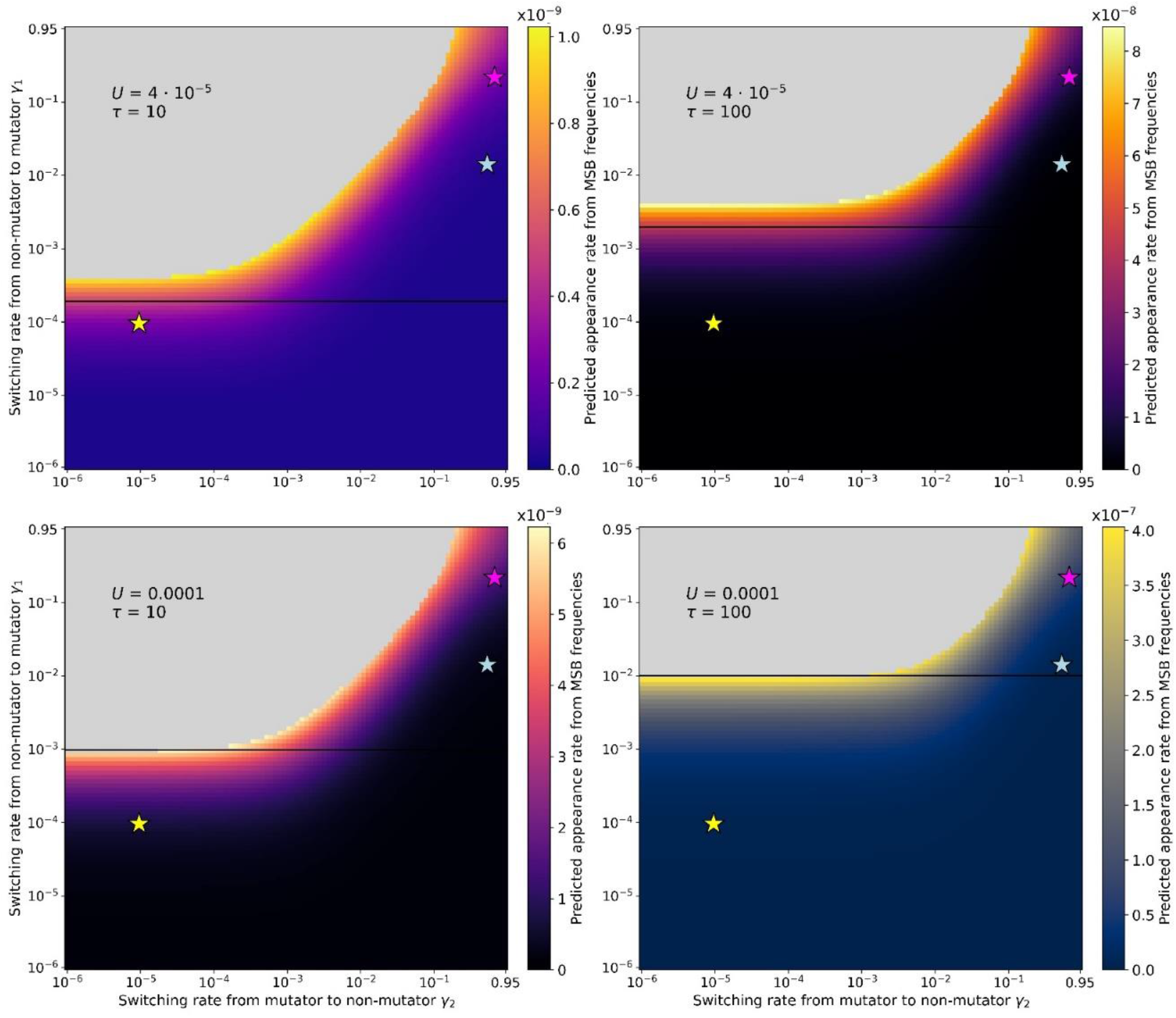
Predicted rate of adaptation for model with gradual transition between the non-mutator to the mutator phenotype and vice-versa. We observe the same patterns as for Figure 2. Parameters: *δ* = 0.01, *s* = 0.03, *β* = 5000.

**Supplementary Figure 11.**
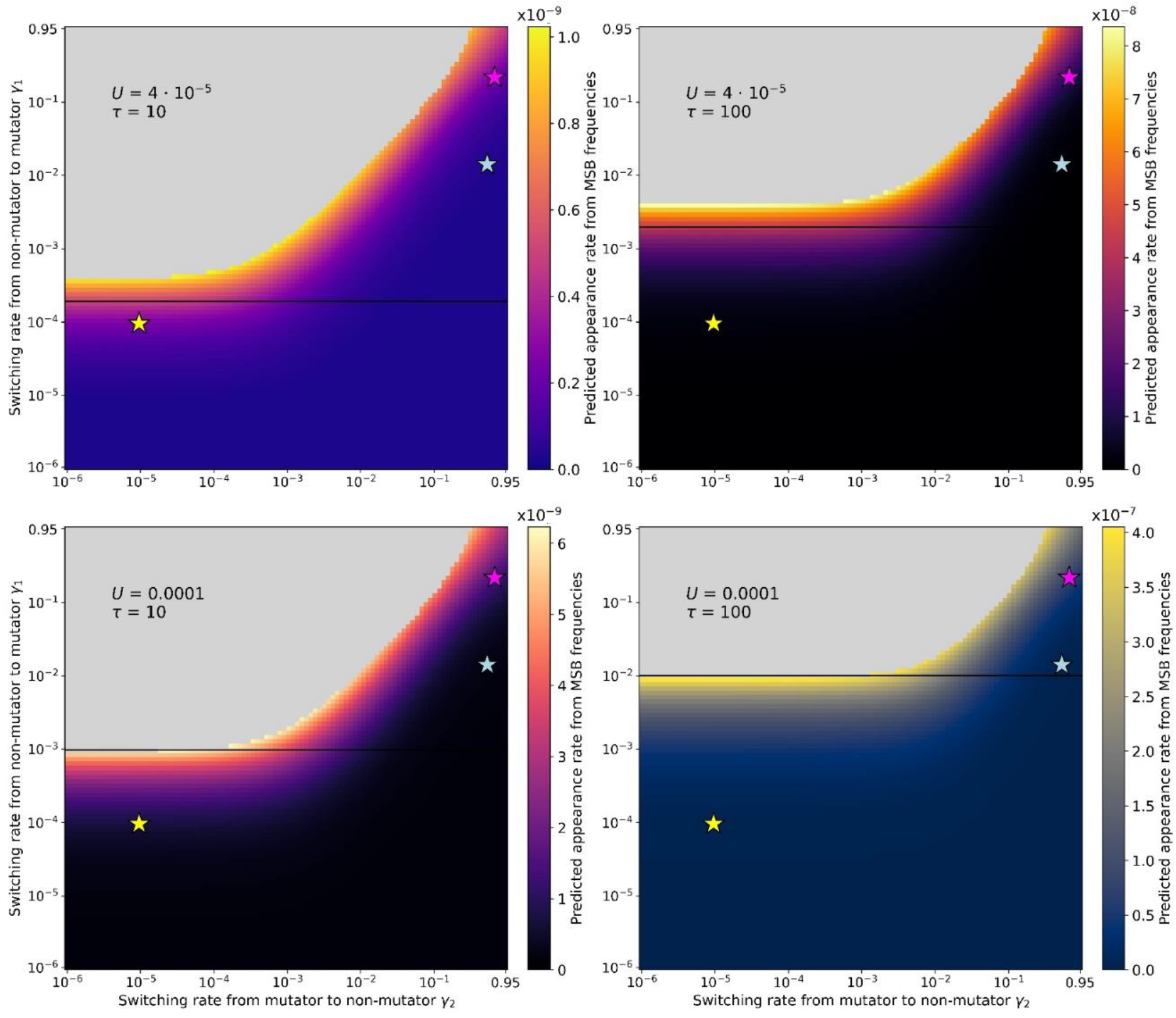
Predicted rate of adaptation for model with gradual transition between the non-mutator to the mutator phenotype and vice-versa. Same as Supplementary Figure 2, except for *δ* = 0.99.

**Supplementary Figure 12.**
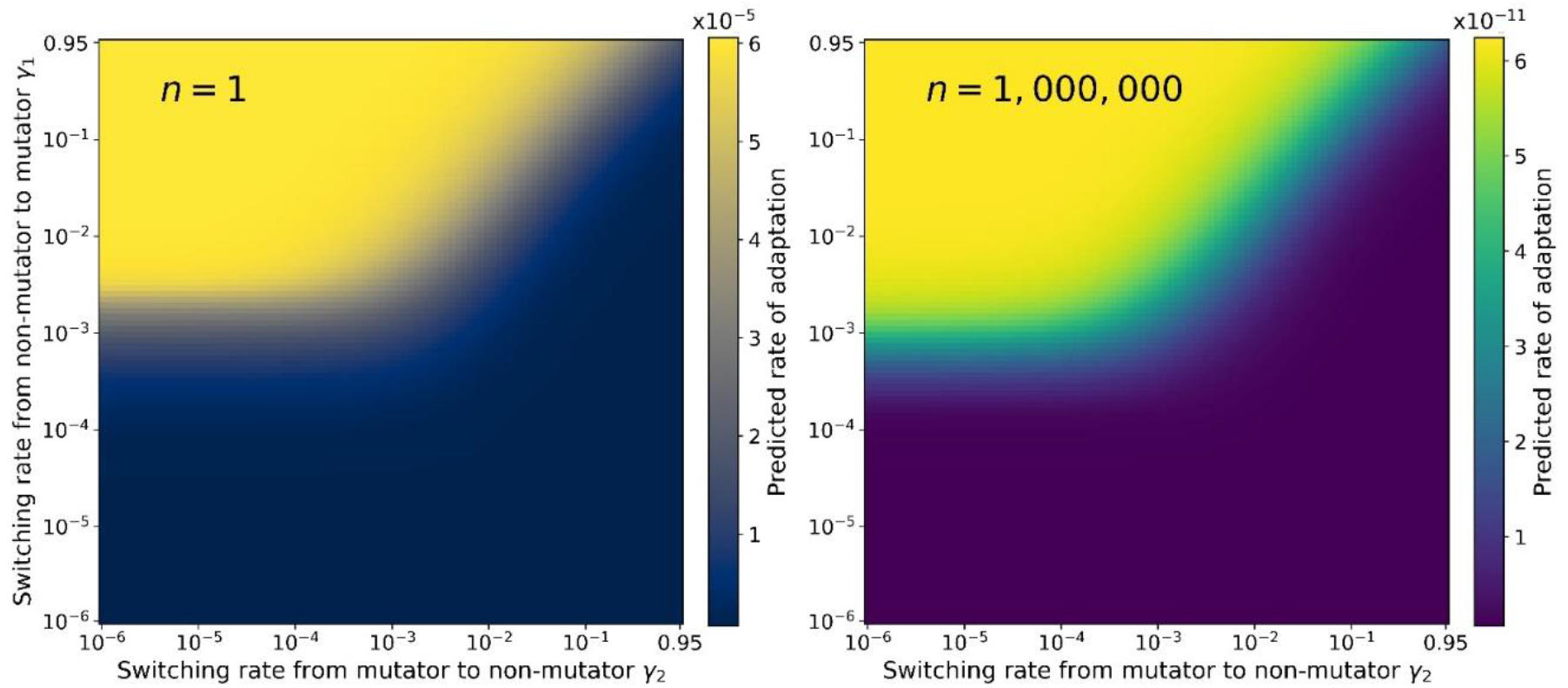
Sensitivity analysis for the number of loci *n*. Although the rate of adaptation is 6 orders of magnitude lower for *n* = 1,000,000, the relative rate of adaptation is similar for the two values of *n*. Indeed, the highest rates of adaptation are observed for 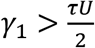 when 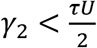 and for *γ*_1_ > *γ*_2_ when 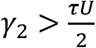. *U* = 0.0001, *s* = 0.03, *τ* = 10.

**Supplementary Figure 13.**
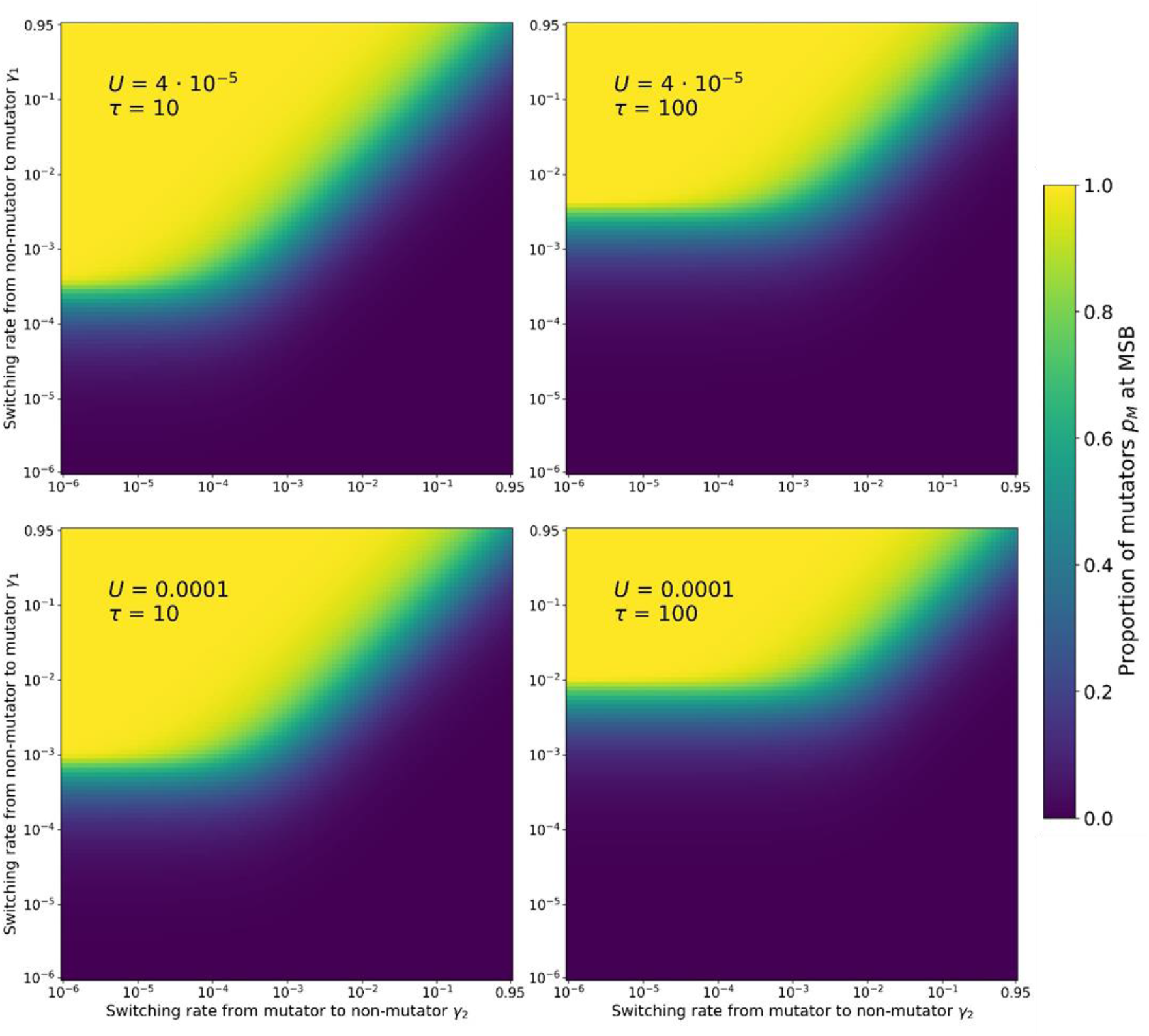
Proportion of mutators *p_M_* at mutation-selection balance. The proportion of mutators increases with the switching rate from non-mutator to mutator. When *γ*_2_ < *τU*/2 and *γ*_1_ > *τU*/2, the proportion of mutators at mutation-selection balance is higher than 0.5, hence excluded from our model. This is also the case for *γ*_2_ > *τU*/2 and *γ*_1_ > *γ*_2_. For *γ*_2_ < *τU*/2, the highest realistic frequencies of mutators are obtained for *γ*_1_ = *τU*/2. For *γ*_2_ > *τU*/2, the highest realistic frequencies of mutators are obtained for *γ*_1_ = *γ*_2_. Parameters: *s* = 0.03, *β* = 5000.

**Supplementary Figure 14.**
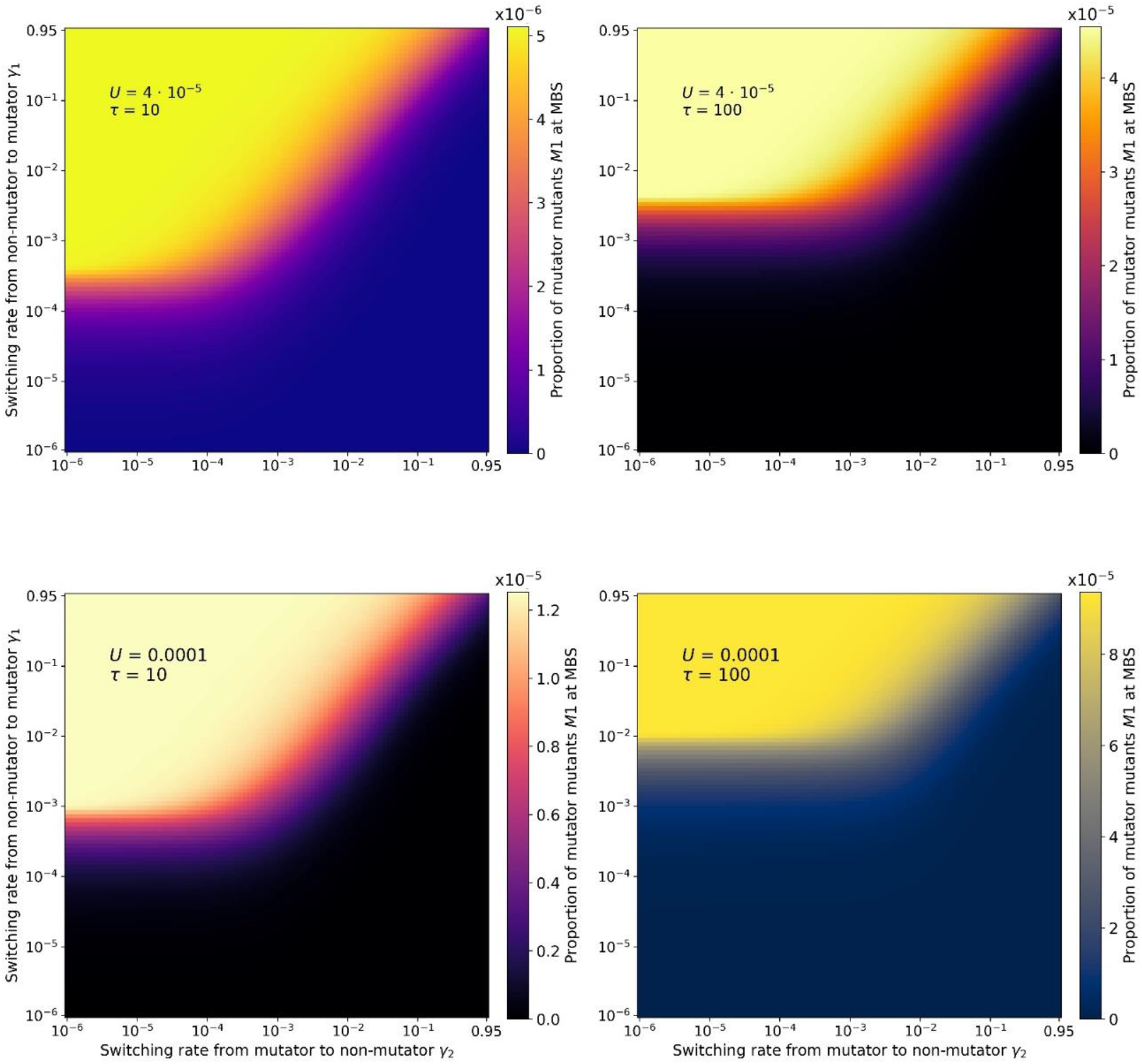
Frequency of mutator mutants *M*_1_ at mutation-selection balance. The proportion of mutator mutants at mutation-selection balance corresponds roughly to the proportion of mutators at mutation-selection balance. However, for negligible inheritance, the frequency of mutator mutants at mutation-selection balance is lower than expected. The frequency of mutator mutants for *γ*_2_ < *τU*/2 and *γ*_1_ = *τU*/2 is higher than expected from the mutator frequency, and is the frequency of mutator mutants for *γ*_2_ > *τU*/2 and *γ*_1_ = *γ*_2_, except when either *γ*_1_ or *γ*_2_ are superior to 0.01. Parameters: *s* = 0.03, *β* = 5000.

**Supplementary Figure 15.**
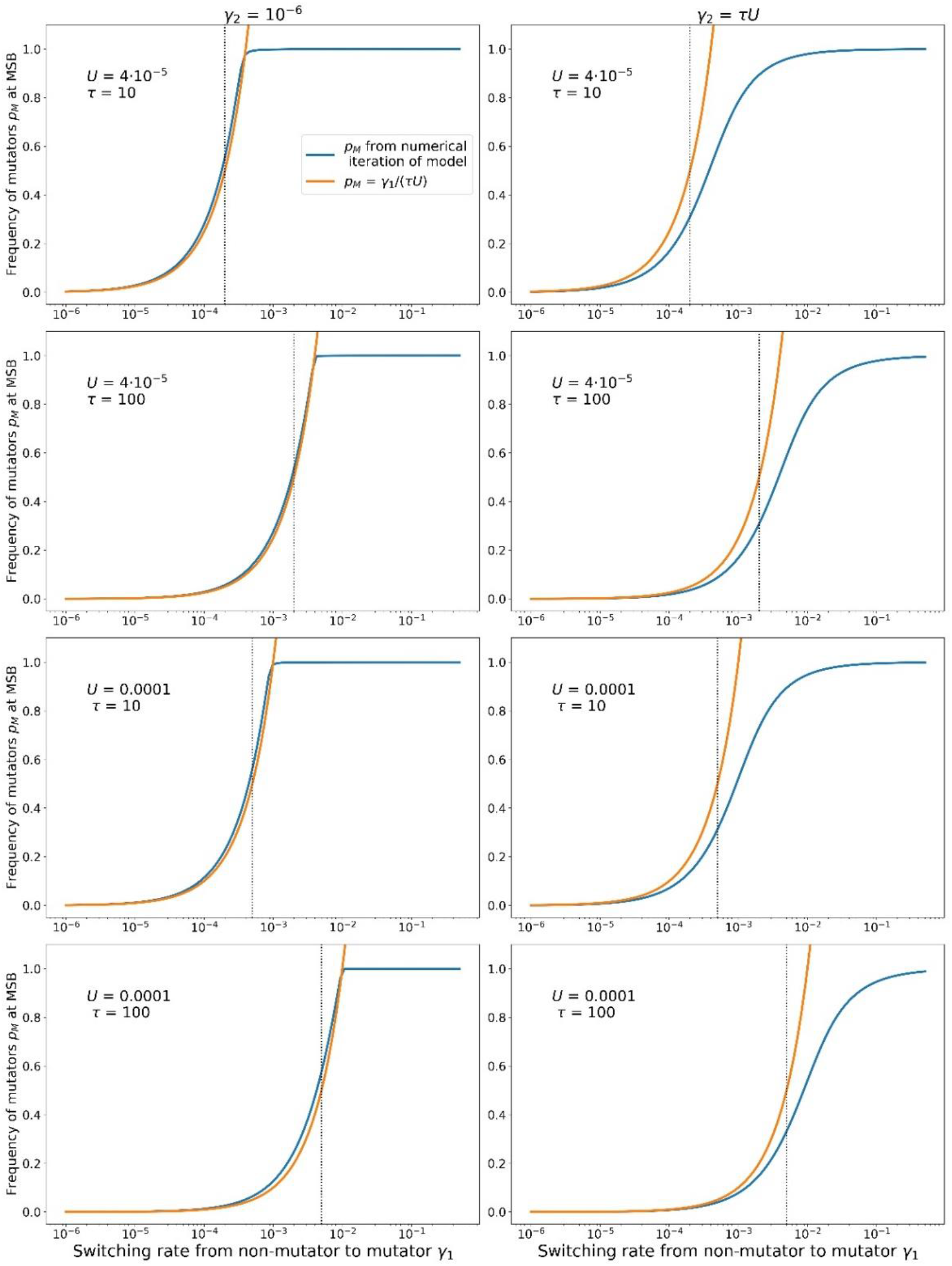
Comparison of the proportion of mutators from the numerical simulation with the analytical prediction derived by [33]. The two considered switching rates from mutator to non-mutator *γ*_2_ correspond to genetic inheritance (*γ*_2_ = 10^−6^) and the highest value of *γ*_2_ for which we use Eq. 4 (*γ*_2_ = *τU*/2). The dashed line corresponds to *γ*_1_ = *τU*/2. For *γ*_2_ = 10^−6^, we observe an excellent fit for all considered parameter sets. For *γ*_2_ = *τU*/2, the fit worsens progressively with increasing switching rate from non-mutator to mutator *γ*_1_. Parameters: *s* = 0.03, *β* = 5000.

**Supplementary Figure 16.**
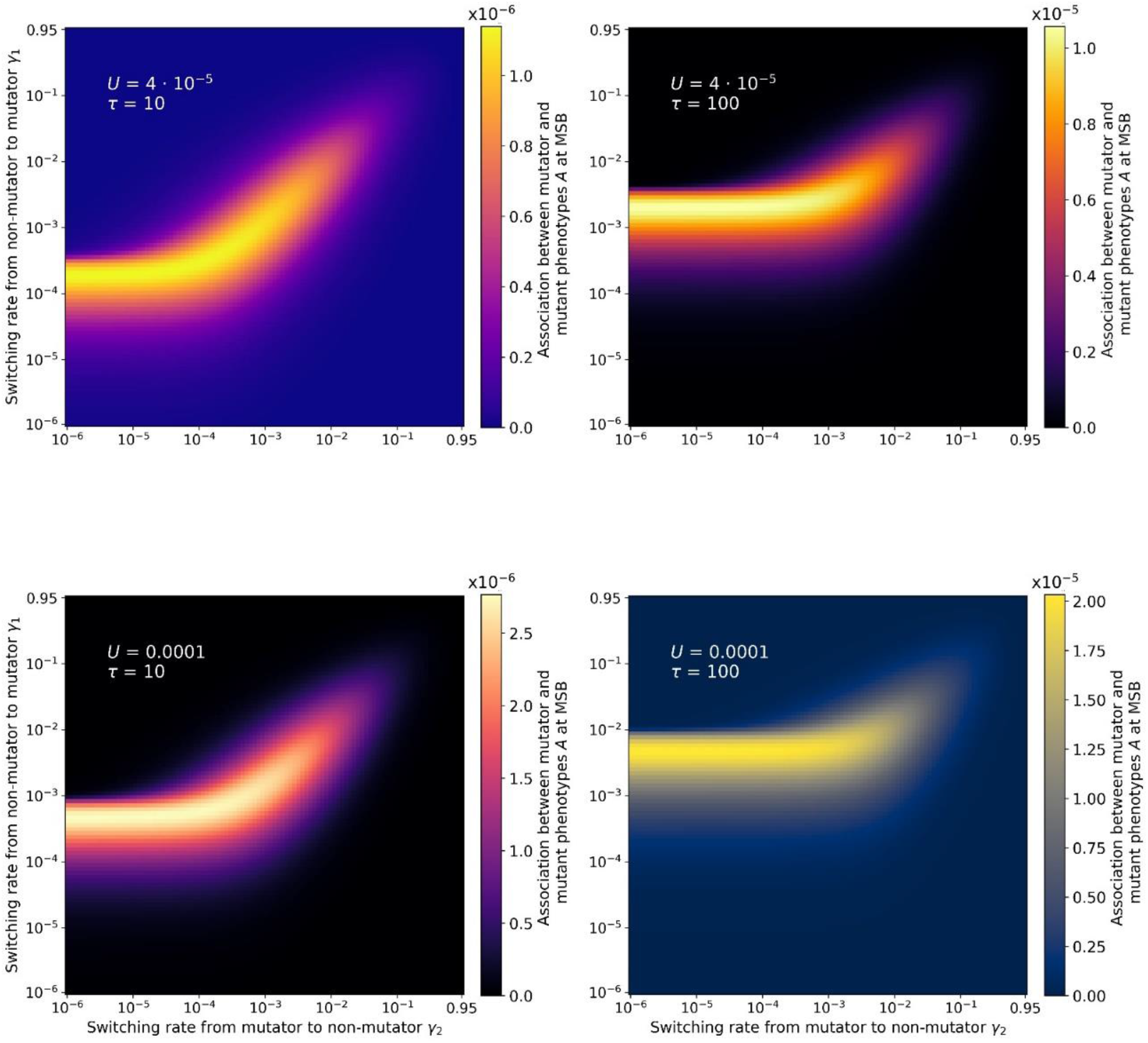
Association between mutator and mutant phenotypes. The plotted value is the difference between the expected frequency of mutator mutants at mutation-selection balance if mutator and mutant phenotypes where independent, and the frequency obtained from the numerical iteration of our model (see Eq. 6). We observe that the regions with highest association correspond to the regions with highest rate of adaptation presented in Figure 2. These regions correspond to *γ*_1_ = *τU*/2 when *γ*_2_ < *τU*/2, and *γ*_1_ = *γ*_2_ when *γ*_2_ > *τU*/2. Parameters: *s* = 0.03, *β* = 5000.

**Supplementary Figure 17.**
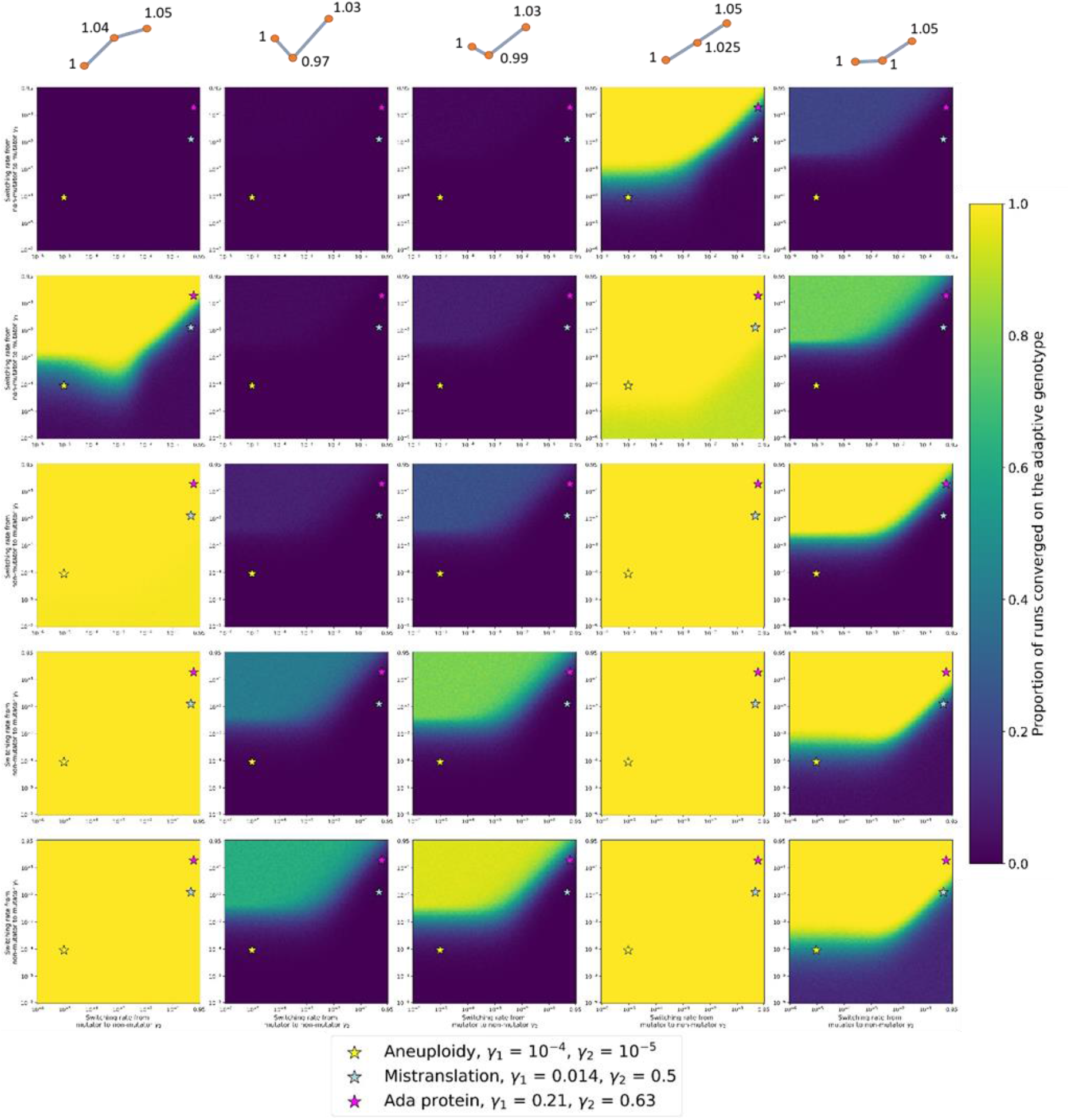
Evolution of the progression of the number of runs that converged to the double mutant, which has the highest fitness. The cartoon representation above each column represents the shape of the fitness landscape on which the stochastic simulation was run. The numbers next to each genotype correspond to the fitness of each genotype relative to the wild-type. The coloured stars represent the three systems of non-genetic inheritance of the mutation rate described in the empirical literature (see legend). Parameters: *U* = 4 · 10^−5^, *τ* = 100, *N* =, *β* = 5000, *s* = 0.03.

**Supplementary Figure 18.**
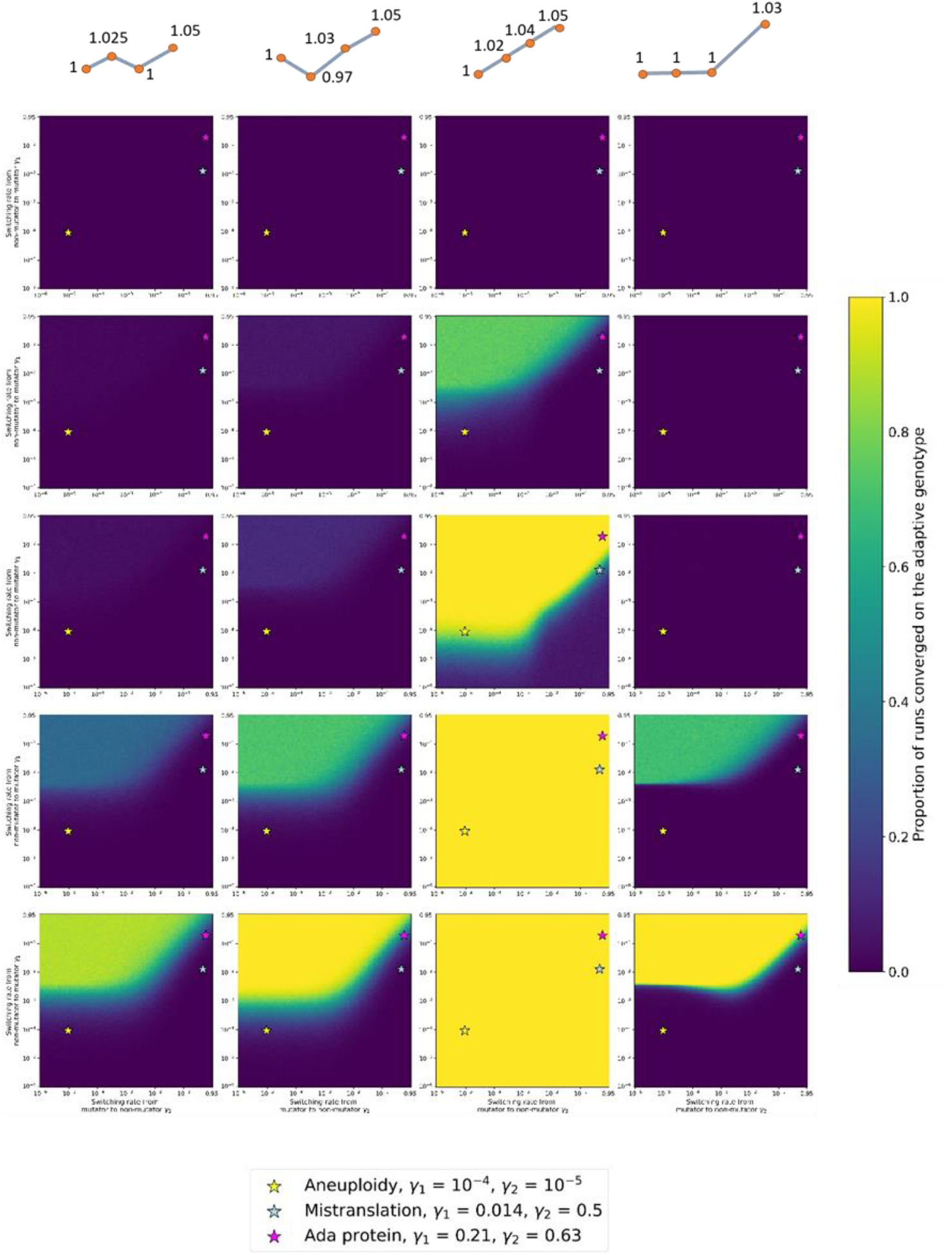
Evolution of the progression of the number of runs that converged to the triple mutant, which has the highest fitness. The cartoon representation above each column represents the shape of the fitness landscape on which the stochastic simulation was run. The numbers next to each genotype correspond to the fitness of each genotype relative to the wild-type. The coloured stars represent the three systems of non-genetic inheritance of the mutation rate described in the empirical literature (see legend). Parameters: *U* = 4 · 10^−5^, *τ* = 100, *N* =, *β* = 5000, *s* = 0.03.

**Supplementary Figure 19.**
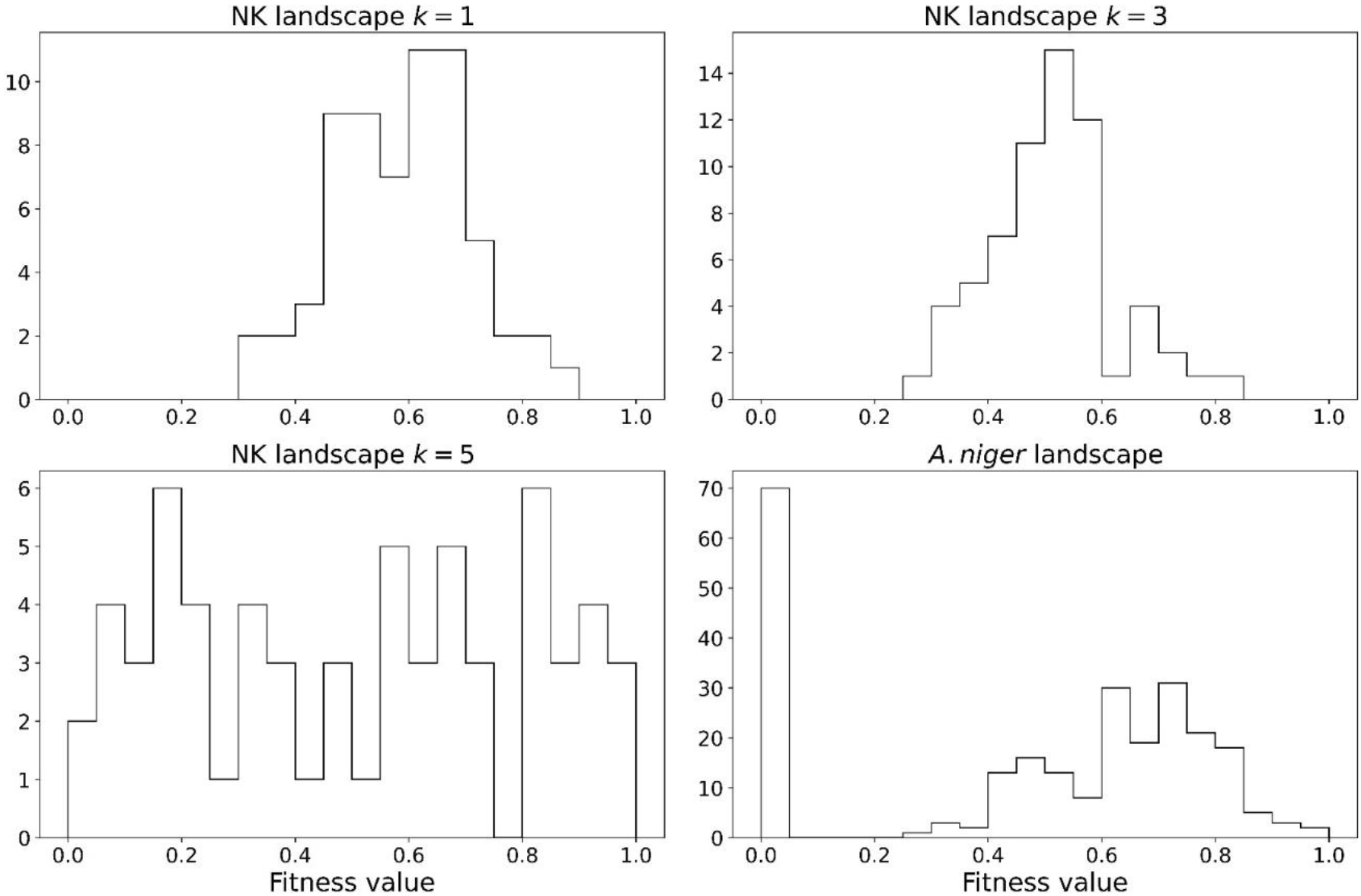
Distribution of fitness values in the NK and *Aspergillus niger* landscapes. In the NK landscapes, the higher the *k* (representing the ruggedness), the wider the distribution of fitness values.

**Supplementary Figure 20.**
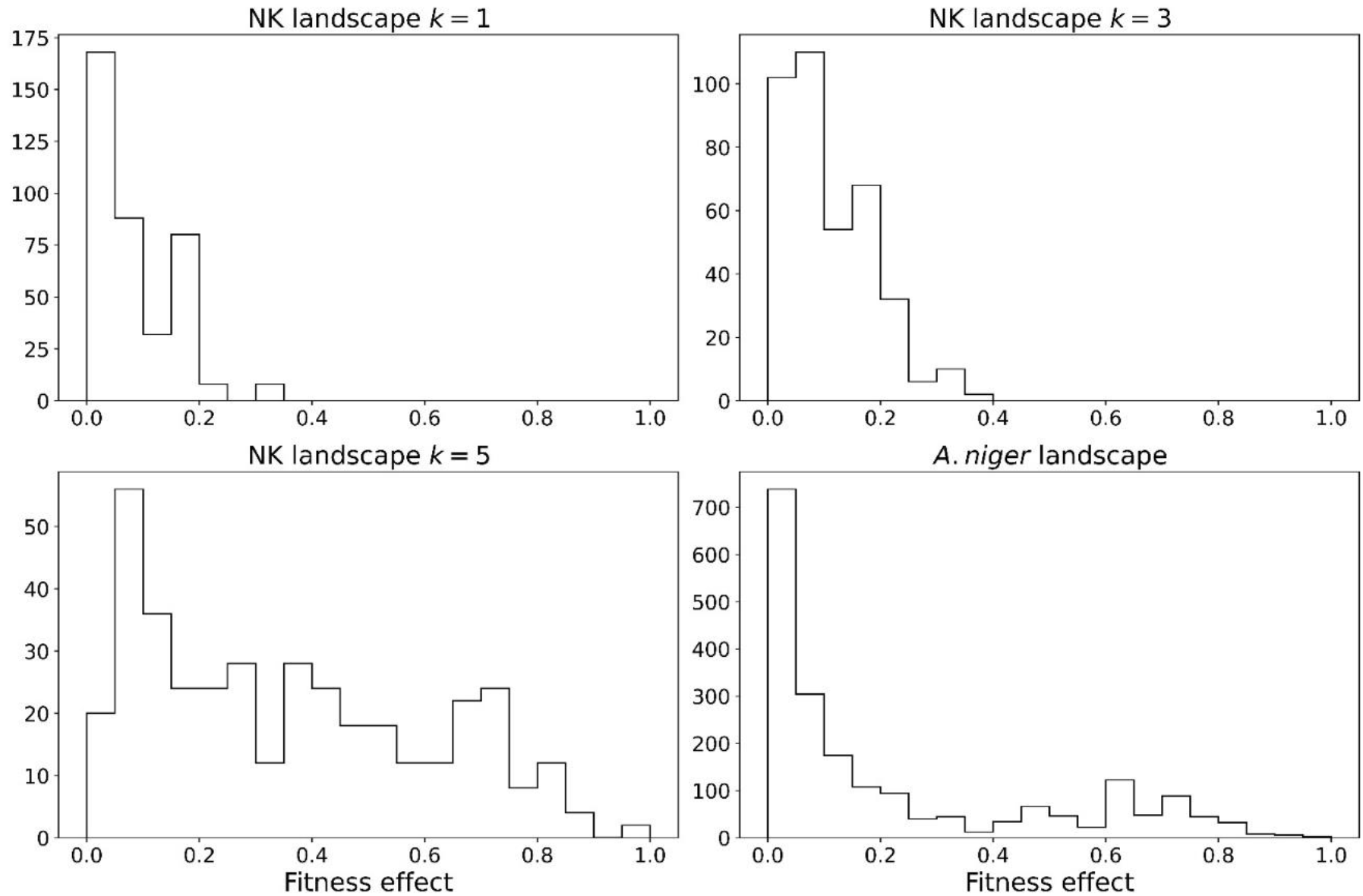
Distribution of fitness effects in the NK and *Aspergillus niger* landscapes. A histogram of the mean difference between each genotype and its single mutants for different values of the ruggedness parameter *k*. As expected, as *k* increases, the distribution is wider.

**Supplementary Figure 21.**
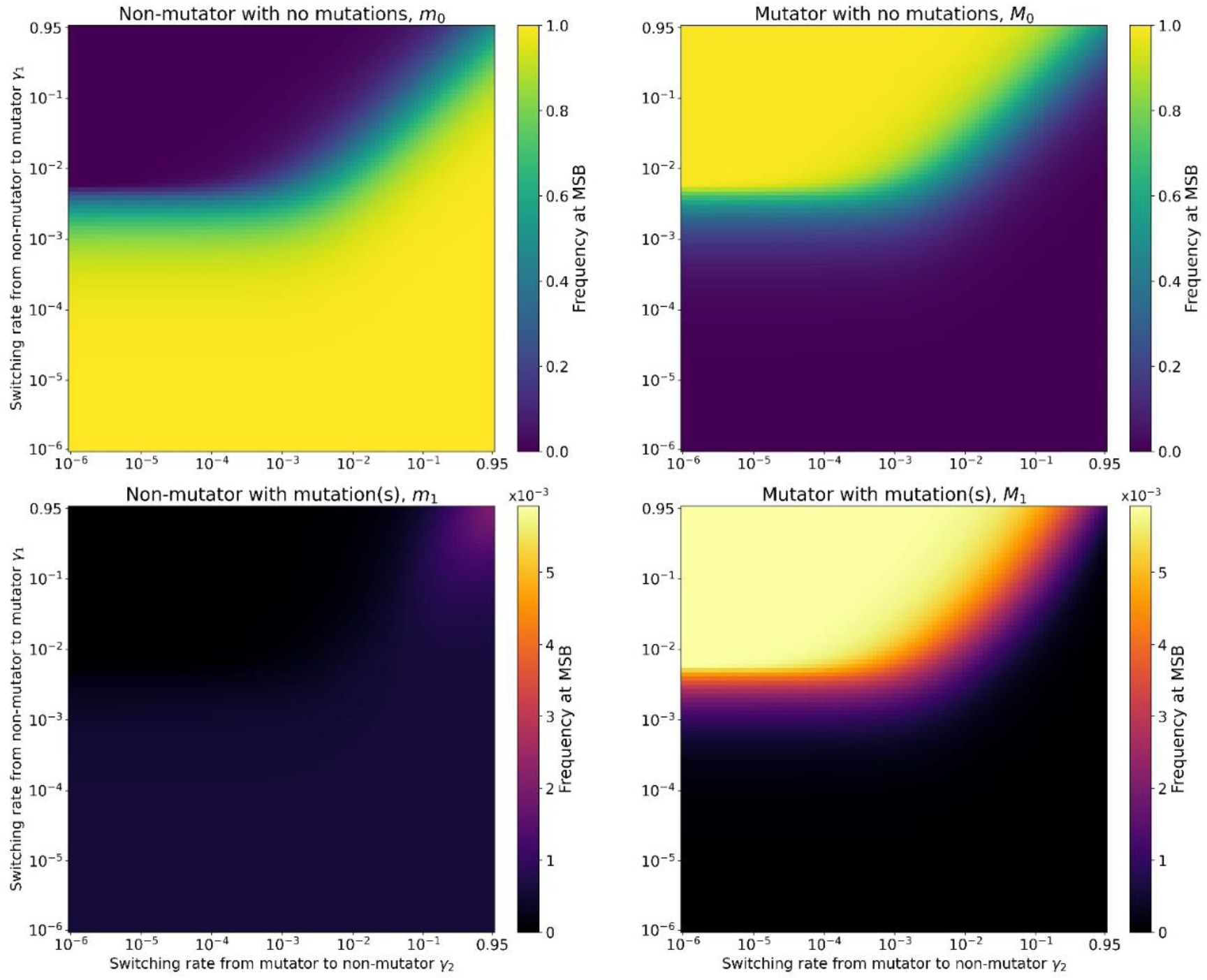
Proportion of non-mutator with no mutations, mutator with no mutations, non-mutator single mutant, and mutator single mutant at mutation-selection balance. Here, computed for the NK landscape *k* = 1. Parameters: *U* = 10^−4^, *τ* = 10.

**Supplementary Figure 22.**
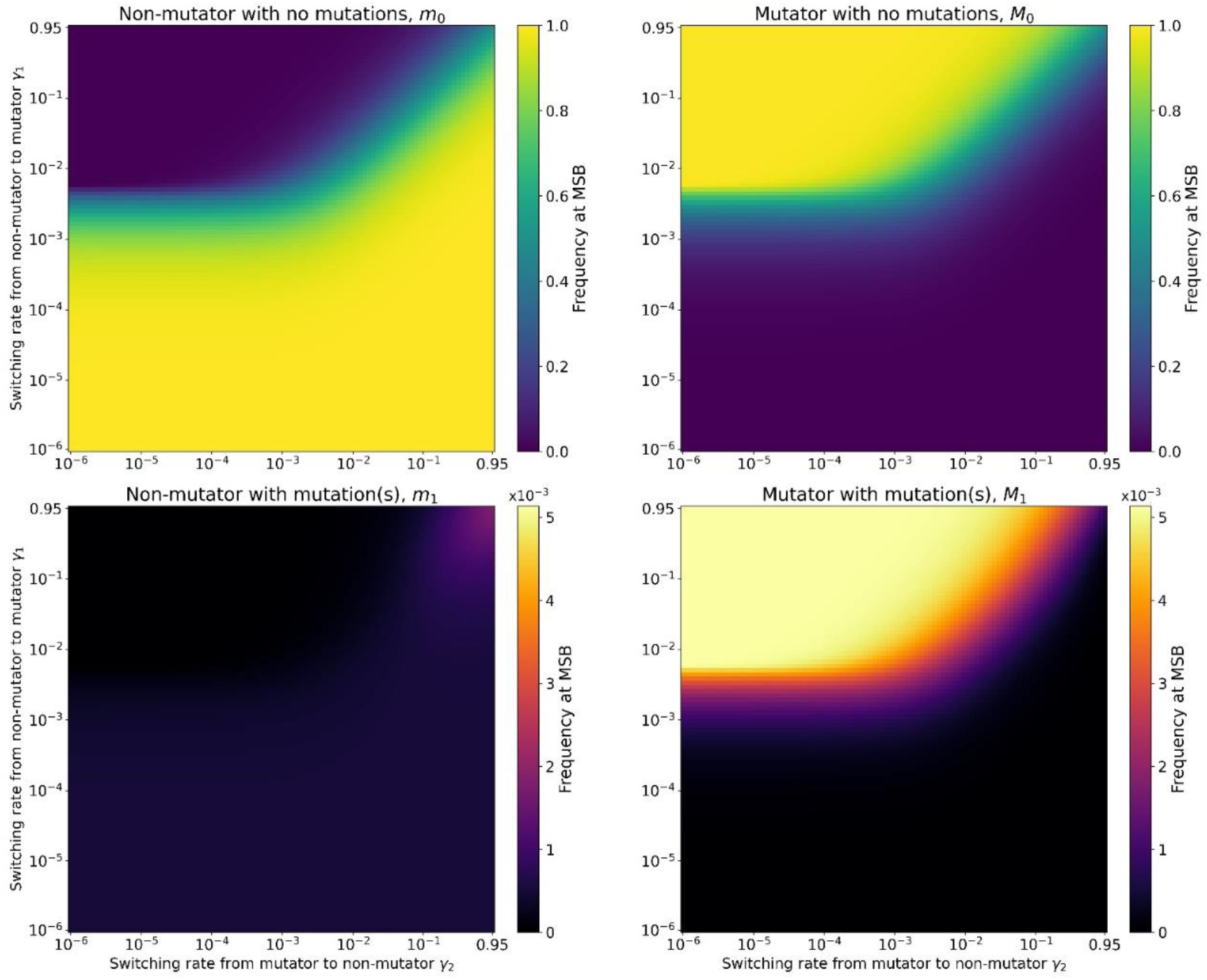
Proportion of non-mutator with no mutations, mutator with no mutations, non-mutator single mutant, and mutator single mutant at mutation-selection balance. Same as Supplementary Figure 21, except for the NK landscape *k* = 3. We observe no substantial difference with Supplementary Figure 21, plotted for the NK landscape *k* = 1. Parameters: *U* = 10^−4^, *τ* = 10.

**Supplementary Figure 23.**
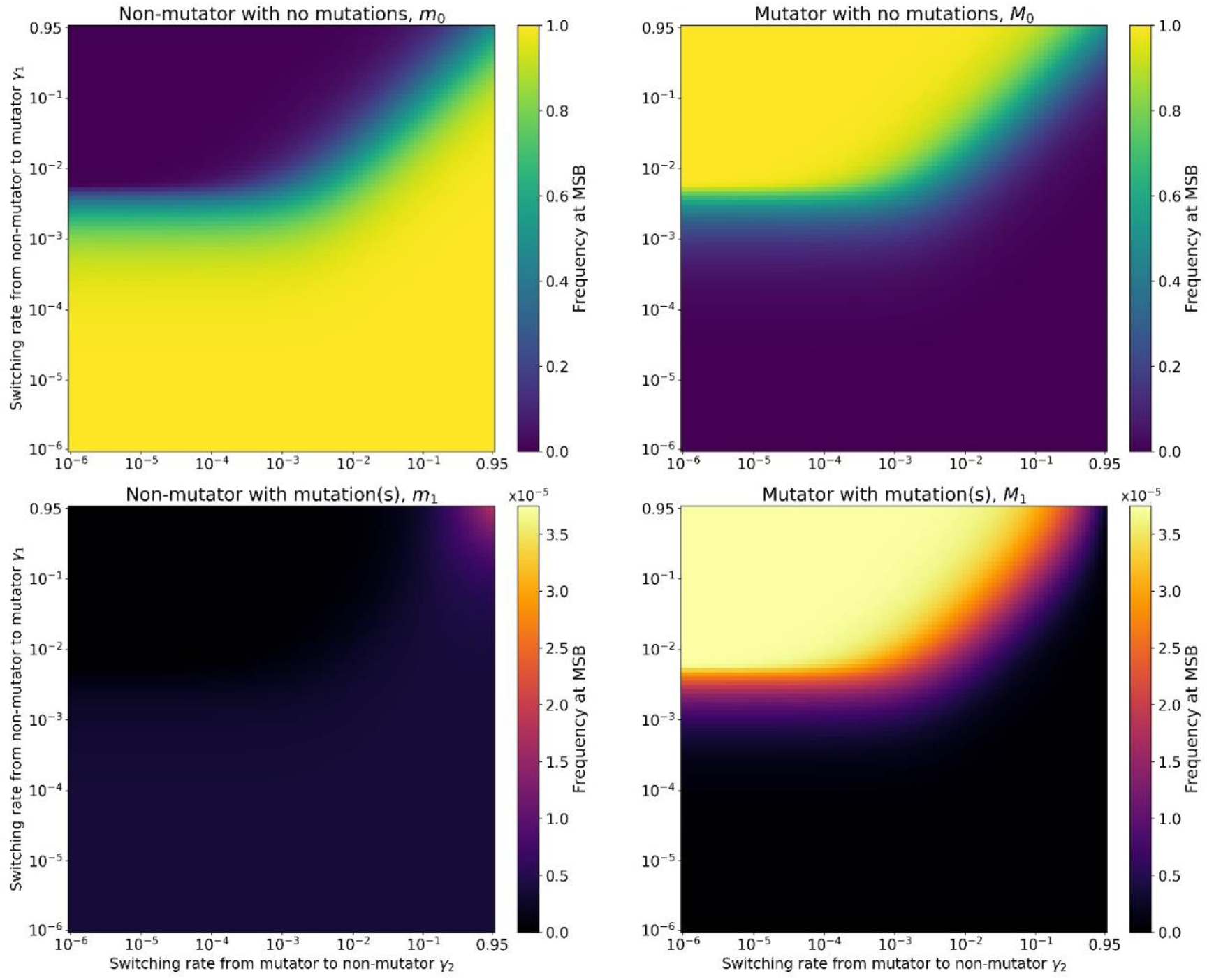
Proportion of non-mutator with no mutations, mutator with no mutations, non-mutator single mutant, and mutator single mutant at mutation-selection balance. Same as Supplementary Figure 21, except for the NK landscape *k* = 5. We observe no substantial difference with Supplementary Figure 21, plotted for the NK landscape *k* = 1. Parameters: *U* = 10^−4^, *τ* = 10.

**Supplementary Figure 24.**
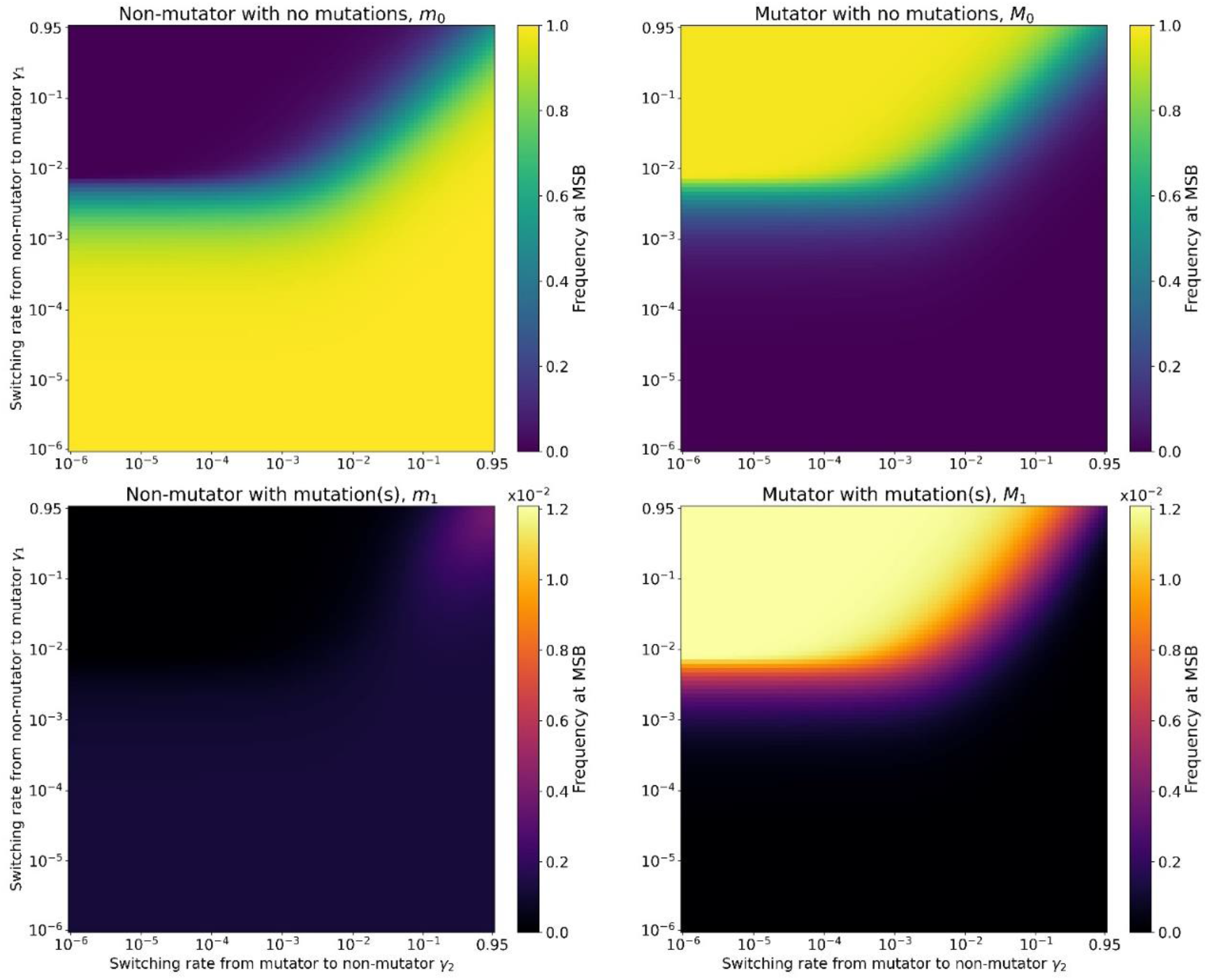
Proportion of non-mutator with no mutations, mutator with no mutations, non-mutator single mutant, and mutator single mutant at mutation-selection balance. Same as Supplementary Figure 21, except for empirical fitness landscape of *Aspergillus niger*. Parameters: *U* = 10^−4^, *τ* = 10.

**Supplementary Figure 25.**
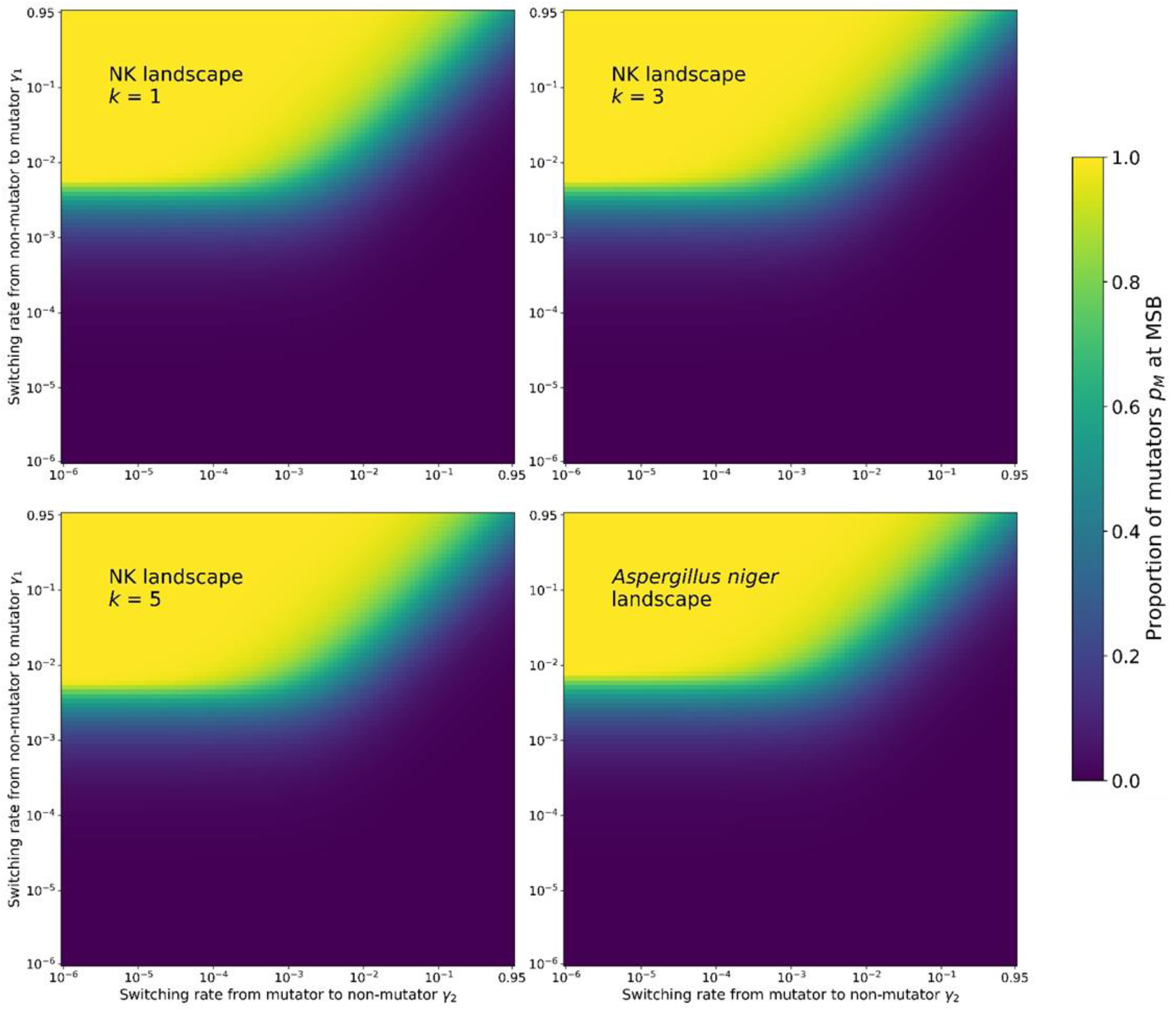
Proportion of mutators *p_M_* at mutation-selection balance for complex landscapes. We observe similar results to the simple landscape, see Supplementary Figure 13, with the mutator frequency being equal to 0.5 for *γ*_1_ = *τU*/2 when *γ*_2_ < *τU*/2 and *γ*_1_ = *γ*_2_ when *γ*_2_ > *τU*/2. Parameters: *U* = 4 · 10^−5^, *τ* = 100, *s* = 0.03.

**Supplementary Figure 26.**
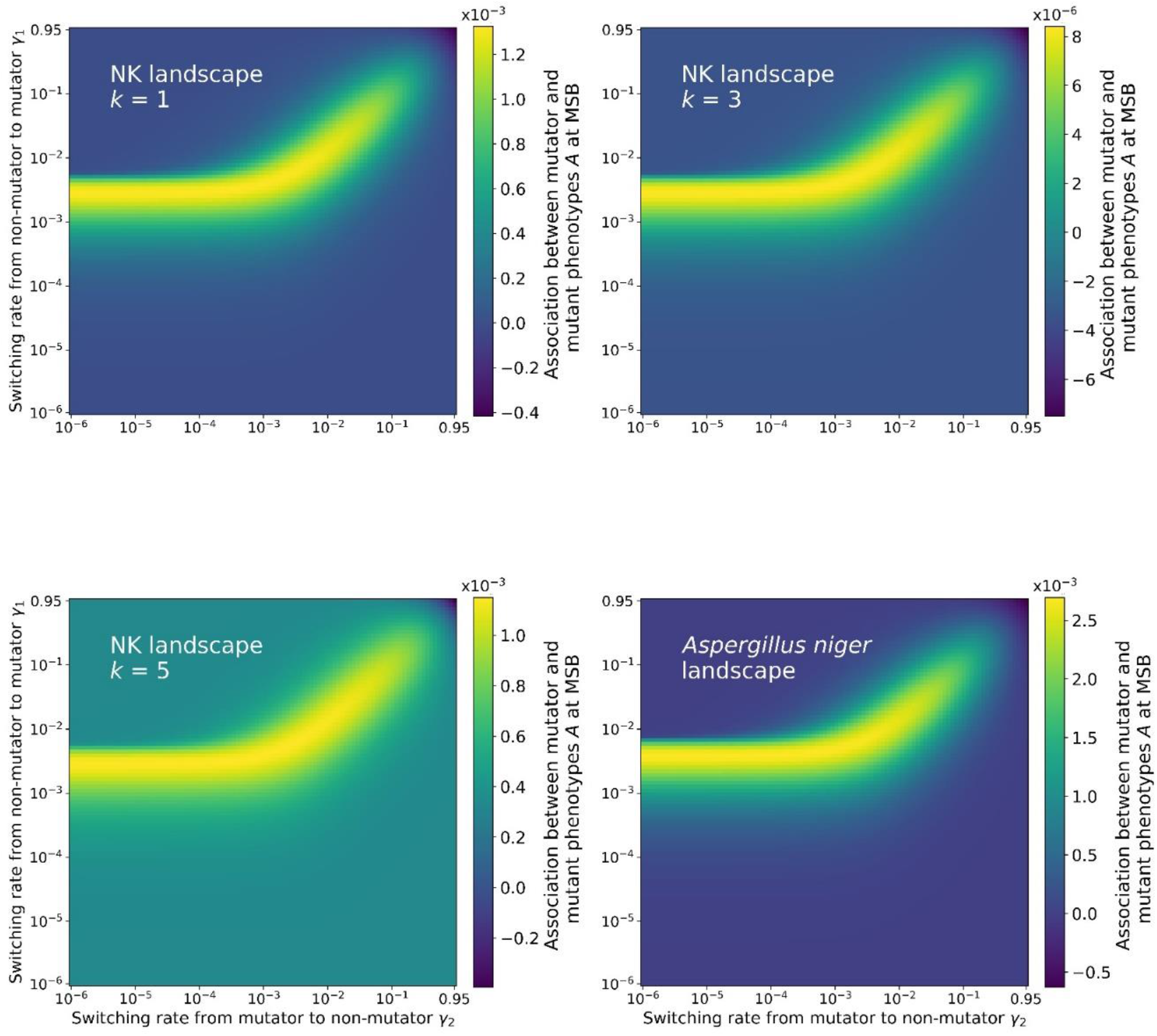
Association between mutator and mutant phenotypes for complex landscapes. The plotted value is the difference between the expected frequency of mutator mutants at mutation-selection balance if mutator and mutant phenotypes where independent, and the frequency obtained from the numerical iteration of our model (see Eq. 6). We observe similar patterns as for the simple landscape in Supplementary Figure 11. Parameters: *U* = 4 · 10^−5^, *τ* = 100, *s* = 0.03.

**Supplementary Figure 27.**
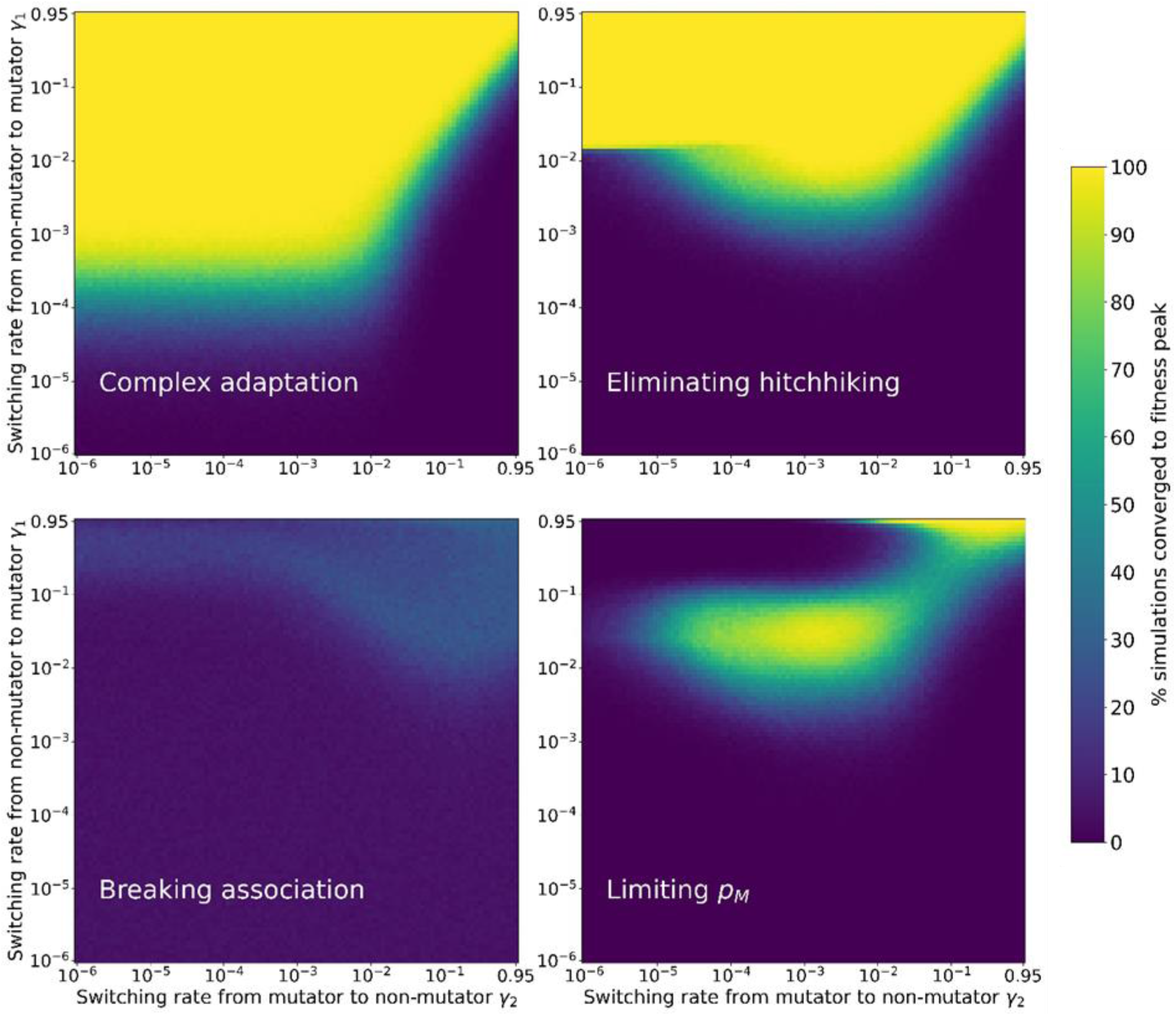
Complex adaptation on NK landscape, *k=1*. The proportion of runs out of 1000 that converged upon the fittest genotype in the landscape was recorded after 500 generations. In order to disentangle the different effects of the non-genetic inheritance of the mutation rate on the rate of adaptation, we then rerun the simulation while removing the association of mutator and mutant, limiting the frequency of mutators during the evolution, and eliminating hitchhiking. Note that hitchhiking is also eliminated when the association between mutator and mutant is broken, or the frequency of the mutator limited. We observe that the region with rates of adaptation for *γ*_1_ < *τU*/2 and *γ*_2_ < *τU*/2 disappears when hitchhiking is eliminated. Breaking association reduces adaptation except for high *γ*_1_ and *γ*_2_. Limiting the frequency *p_M_* reduces adaptation overall. Parameters: *U* = 4 · 10^−5^, *τ* = 100, *N* = 1000.

**Supplementary Figure 28.**
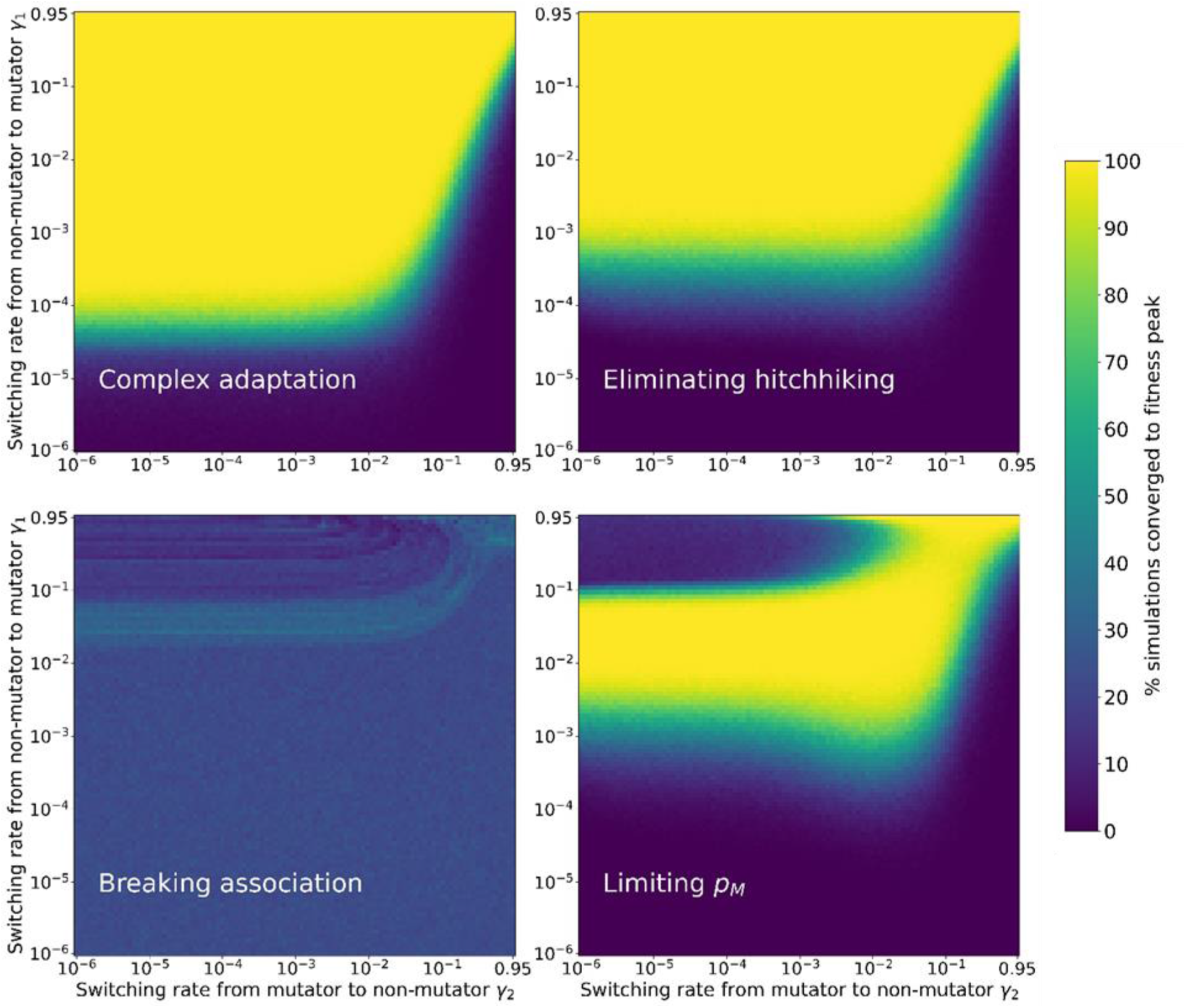
Complex adaptation on NK landscape, *k=3*. Same as Supplementary Figure 27, but for *k=3*. Parameters: *U* = 4 · 10^−5^, *τ* = 100, *N* = 10^7^.

**Supplementary Figure 29.**
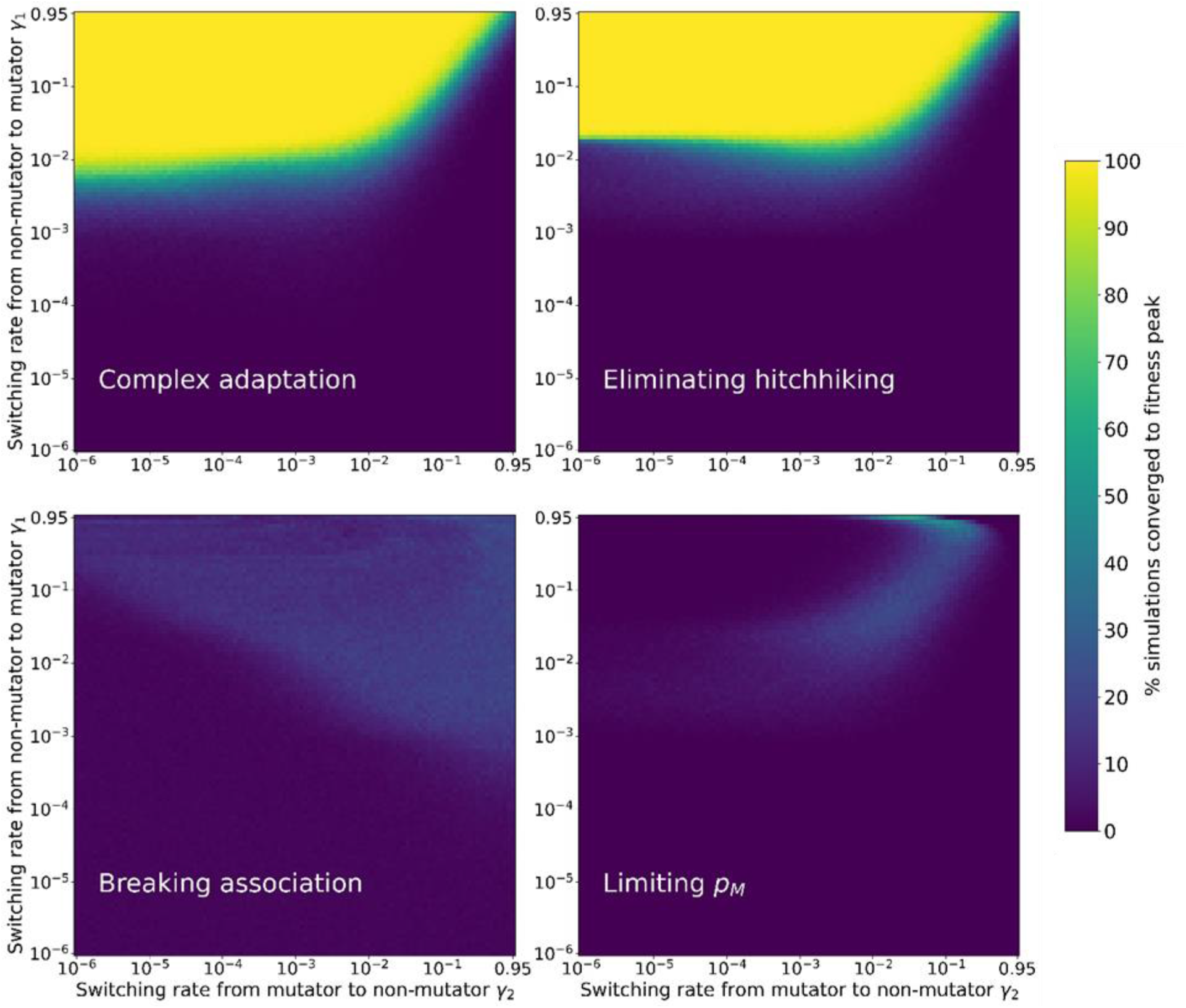
Complex adaptation on NK landscape, *k=5*. Same as Supplementary Figure 27, but for *k=5*. Parameters: *U* = 4 · 10^−5^, *τ* = 100, *N* = 10^7^.

**Supplementary Figure 30.**
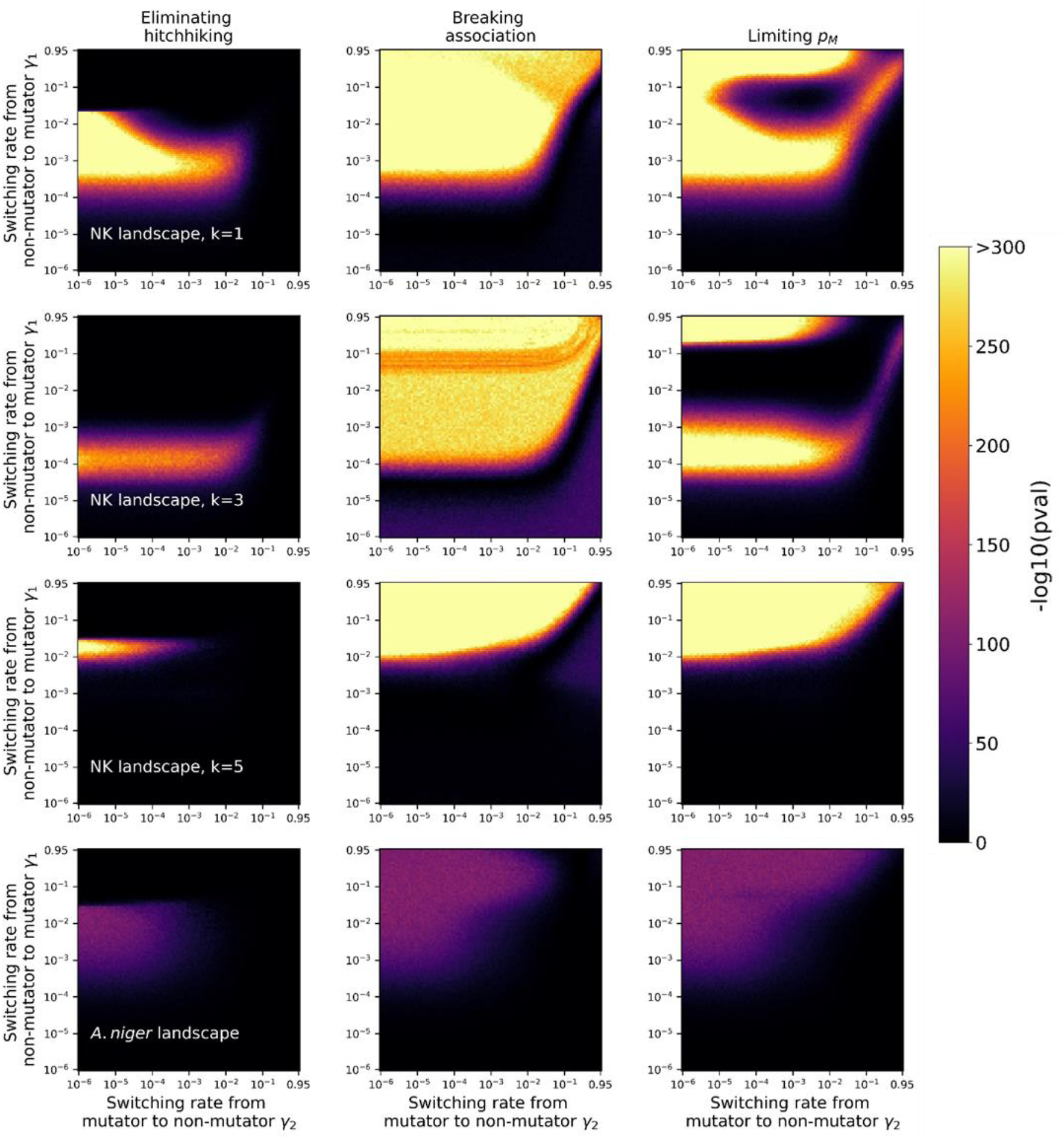
Statistical analysis of the rate of adaptation in the complex adaptation simulation and its modifications: eliminating hitchhiking, reducing the proportion of mutators and breaking association between mutators and mutants. A two-proportion Z-test was performed between the proportion of runs that converged on the adaptive genotype in the complex adaptation simulation and between the proportion of runs that converged on the adaptive genotype in a modification of the simulation. We report the −log10 of the obtained p-value.

